# Conservation of the direct and indirect pathways dichotomy in mouse caudal striatum with uneven distribution of dopamine receptor D1- and D2-expressing neurons

**DOI:** 10.1101/2021.11.04.467262

**Authors:** Kumiko Ogata, Fuko Kadono, Yasuharu Hirai, Ken-ichi Inoue, Masahiko Takada, Fuyuki Karube, Fumino Fujiyama

## Abstract

The striatum is one of the key nuclei for adequate control of voluntary behaviors and reinforcement learning. Two striatal projection neuron types, expressing either dopamine receptor D1 (D1R) or dopamine receptor D2 (D2R) constitute two independent output routes: the direct or indirect pathways, respectively. These pathways co-work in balance to achieve coordinated behavior. Two projection neuron types are equivalently intermingled in most striatal space. However, recent studies revealed two atypical zones in the caudal striatum: the zone in which D1R-neurons are the minor population (D1R-poor zone) and that in which D2R-neurons are the minority (D2R-poor zone). It remains obscure as to whether these imbalanced zones have similar properties on axonal projections and electrophysiology compared to other striatal regions. Based on morphological experiments in mice using immunofluorescence, *in situ* hybridization, and neural tracing, here, we revealed that the poor zones densely projected to the globus pallidus and substantia nigra pars lateralis, with a few collaterals in substantia nigra pars reticulata and compacta. Similar to that in other striatal regions, D1R-neurons were the direct pathway neurons. We also showed membrane properties of projection neurons in the poor zones were largely similar to those in the conventional striatum using *in vitro* electrophysiological recording. In addition, the poor zones existed irrespective of the age or sex of mice. We also identified the poor zones in the common marmoset as well as other rodents. These results suggest that the poor zones in the caudal striatum follow the conventional projection patterns irrespective of imbalanced distribution of projection neurons. The poor zones could be an innate structure and common in mammals. The unique striatal zones possessing highly restricted projections could relate to functions different from those of motor-related striatum.

## 1 Introduction

The striatum regulates voluntary movement and reward-related learning by integrating excitatory inputs from the cerebral cortex and thalamus (Alexander et al., 1986; Hikosaka et al., 2000; Kreitzer and Malenka, 2008; Peak et al., 2019; Redgrave et al., 2011). The medium spiny projection neurons (MSNs)—i.e., the major population of striatal neurons— are classified into two groups: direct and indirect pathway neurons, depending on their projection targets and gene expression. The direct pathway MSNs (dMSNs) directly transmit information to output nuclei, such as the globus pallidus internal segment (GPi, the counter part of the entopeduncular nucleus (EP) in rodents) and substantia nigra (SN). The dMSNs express GABA, dopamine receptor D1 (D1R), and substance P. In contrast, indirect pathway MSNs (iMSNs) express GABA, dopamine receptor D2 (D2R), and enkephalin and project to the globus pallidus external segment (GP in rodents). In turn, GP projects to the output nuclei, namely, iMSNs indirectly project to the output nuclei (Albin et al., 1989; Alexander and Crutcher 1990; Graybiel, 1990). It has long been believed that both types of projection neurons are randomly distributed (Gerfen, 1989; Lança et al., 1986; Tinterri et al., 2018), and each local striatal area contains an almost equal proportion of both types (Hedreen and DeLong, 1991; Selemon and Goldman-Rakic, 1990).

Dense and topographic corticostriatal innervation recruits striatal subregions for specific functions (Nambu, 2008; Shepherd, 2013; Shipp, 2016). The caudal striatum (cStr) in rodents lies under the temporal cortical area as being similar to the caudate tail (CDt) in primates, which is a curved long extension of the ventral part of the caudate nucleus. Temporal areas, including somatosensory, visual, and auditory related areas, innervate cStr in rodents (Deniau et al., 1996; Foster et al., 2021; Hintiryan et al., 2016; Hunnicutt et al., 2016; Jiang and Kim, 2018; Xiong et al., 2015) and CDt in primates (Brown et al., 1995; Caan et al., 1984; Saint-Cyr et al., 1990; Yeterian and Pandya, 1998; Yeterian and Van Hoesen, 1978); therefore, they are considered the sensory striatum. In addition, similar to cStr in rodents (Menegas et al., 2015; Watabe-Uchida et al., 2012), CDt in primates receives the projection from a specific group of dopaminergic neurons (Kim et al., 2014; Kim and Hikosaka, 2013). Thus, cStr in rodents and CDt in primates share common neural connection features. The functional significance of CDt and cStr has been also gradually uncovered. CDt in primates is involved in distinct functions, such as coding value of objects (Griggs et al., 2017; Kim et al., 2017; Kim et al., 2014; Kim and Hikosaka, 2013). Recent studies in rodents have shown that the cStr is involved in the avoidance behavior of mice (Menegas et al., 2018; Menegas et al., 2017).

Gangarossa et al. (2013b) revealed a unique cStr region adjacent to GP using BAC transgenic mice that express eGFP. The region is surprisingly composed almost exclusively of *Drd1a* expressing neurons and is therefore called D2R/A2aR-expressing MSNs-poor zone (Gangarossa et al., 2013b). Studies with transgenic mice also showed the area with fewer D1R-expressing neurons in the cStr (Miyamoto et al., 2018; Miyamoto et al., 2019; for review, see Valjent and Gangarossa, 2021). Our previous study with wild-type mice also confirmed the highly uneven distribution of D1R and D2R immunoreaction in the unique regions of cStr (Ogata et al., 2018).

Although such uneven distribution of D1R and D2R immunoreactivity conjures up the possibility that the direct and indirect pathway neurons are separately distributed in the unique region of cStr, it contradicts the conservative model according to which the physiological function requires a balance of the direct and indirect pathway neurons mediated by D1R and D2R, respectively (Calabresi et al., 2014; Cui et al., 2013; Friend and Kravitz, 2014; Isomura et al., 2013). Thus, we raise two questions: First, does the uneven distribution of D1R and D2R immunoreactions actually reflect separate distribution of the direct and indirect pathway projection neurons? If so, what are neural circuitries driven by? Second, is this uneven distribution a common property of the sensory related striatum and conserved among rodents and primates, or is it a rodent-specific feature (Gangarossa et al., 2019)? If the former is true, sensory inputs may modify the uneven structure of the sensory related striatum. Thus, the uneven structure may vary developmentally, since neonatal mice do not respond to visual and auditory stimuli (Huberman et al., 2008; Sonntag et al., 2009), while aged C57BL6 mice typically have impaired hearing (Zheng et al., 1999). Alternatively, the uneven structure of the cStr can be innate, although maturation of MSNs continues in the early postnatal days (Krajeski et al., 2019). To address these questions, we employed a combination of *in situ* hybridization, immunohistochemistry, electrophysiological recording, and retrograde/anterograde tracing in mice, and also compared the cStr structure across ages and species.

## 2 Materials and Methods

All animal experiments in mice and rats were approved by and performed in accordance with the guidelines for the care and use of laboratory animals established by the Committee for Animal Care and Use of Doshisha University (Approval number: A16008, A17001, A18001, A19036, and A20057) and the Animal Care and Use Committee of Hokkaido University (Approval Number: 20- 0106). All animal studies in the common marmosets were conducted in accordance with experimental procedure protocols approved by the Animal Welfare and Animal Care Committee of the Primate Research Institute of Kyoto University (Approval number: 2017-031). All efforts were made to minimize animal suffering and the number of animals used. Chemicals were purchased from Nacalai Tesque (Kyoto, Japan) and Wako (Osaka, Japan), unless otherwise noted.

In this study, 42 wild type C57BL/6J male mice (8-day-old to 172-week-old), three wild type slc:ICR male mice (11-week-old), three Long-Evans male rats (11—13-week-old), three Wistar male rats (12-week-old), and two common marmosets (a 6.8-year-old male and a 5.2-year-old female) were utilized. Additionally, three female BL6 mice were also used to examine sexual differences (10–12-week-old).

### 2.1 Animal surgery: Retro- and anterograde tracing study

Mice were anesthetized by inhalation of isoflurane (Pfizer Japan Inc., Tokyo, Japan) followed by intramuscular injection of a mixture of ketamine (Ketalar; Daiichi-Sankyo, Tokyo, Japan; 40 mg/kg) and xylazine (Bayer HealthCare, Tokyo, Japan; 4 mg/kg). Before and after surgery, butorphanol solution (Meiji Seika Pharma Co Ltd, Tokyo, Japan) was injected subcutaneously (0.2 mg/kg) for analgesia. Each mouse was then fixed to a stereotaxic device (Narishige, Tokyo, Japan). During surgery, the body temperature of the mice was monitored and maintained at approximately 38 ℃ (BWT 100A animal warmer; Bio Research Center, Nagoya, Japan). The skull was drilled into to make a small hole at an appropriate position in accordance with the mouse brain atlas (Paxinos and Franklin, 2013). For retrograde labeling of projection neurons in the rostral and caudal striatum simultaneously, a large volume (> 0.5 µL) of 0.2% cholera toxin subunit B – Alexa Fluor 555 (CTB555) or 488 (CTB488) conjugate (C22843 or C-22841, Thermo fisher Scientific, Inc.) in 0.1 M phosphate buffer (PB, pH 7.4) was injected around the output nuclei of the basal ganglia of 3 mice [anteroposterior (AP): 2.8 mm caudal from the bregma (AP −2.8), lateromedial: 2.1 mm lateral from the midline (L2.1), depth: 3.7—4.2 mm from the pial surface (D 3.7—4.2)] using a glass pipette (tip diameter, 15—20 µm) through which air pressure pulses were delivered with a pressure injector (PV820, World Precision Instruments, Sarasota, FL, USA). The site of injection extended to the substantia nigra pars reticulata (SNpr), substantia nigra pars compacta (SNpc), and EP, but did not reach the GP. For retrograde tracing from substantia nigra pars lateralis (SNpl) or medial geniculate nucleus (MG), 0.2% CTB555 (∼0.2 µl) was injected by pressure, or 5% fluorogold (FG) dissolved in phosphate-buffered saline (PBS, pH 7.4) was injected iontophoretically (0.8—1.0 μA positive current pulses with a 7 s-on/off cycle for 5 min using A365, World Precision Instruments). The coordinates of the injection were [AP −2.9, L 1.8, D 3.8] for SNpl and [AP −3.2, L 2.0, D 2.8] for MG (N = 3 mice for each).

For anterograde tracing, 10% biotinylated dextran amine (BDA 10 kD; D1956, Invitrogen) in PBS, or 2.5% Phaseolus Vulgaris Leucoagglutinin (PHAL; L-1110, Vector laboratories) in 10 mM phosphate (pH 8.0) was injected in the cStr using a glass pipette (tip diameter, 15—25 µm). The BDA solution was injected iontophoretically by a 1.2 μA current pulses with a 7 s-on/off cycle for 5—20 min, or ejected by a single air pulse. PHAL was injected by 4 μA current pulses with a 7 s-on/off cycle for 5 min. After 3 — 5 days, each mouse was perfused as described below. The targeted coordinates for the caudo-dorsal (cdStr) were [AP −1.2, L 2.7, D 1.7], and those for D1R- and/or D2R-poor zones were [AP −1.2, L 2.7, D 2.8] as described in the mouse brain atlas (Paxinos and Franklin, 2013).

### 2.2 Immunofluorescence labeling and tracer visualization

#### 2.2.1 Tissue preparation

Mice and rats were deeply anesthetized with isoflurane and sodium pentobarbital (100 mg/kg, i.p.; Kyoritsu Seiyaku Corporation, Tokyo, Japan). The animals were then transcardially perfused with 8.5% sucrose in 20 mM PB containing 5 mM MgCl2, followed by 4% w/v paraformaldehyde and 75% saturated picric acid in 0.1 M PB. After the perfusion pump was switched off, the brain was postfixed *in situ* for 1.5 h at room temperature (RT; 25 ± 3 ℃), and then the brain was removed from the skull, followed by cryoprotection with 30% sucrose in PBS for 24—48 h at 4 ℃. Tissue blocks containing the striatum were sectioned sagittally or coronally using a freezing microtome (Leica Microsystems, Wetzlar, Germany) at a thickness of 20 µm. Six series of floating sections were collected in 0.1 M PB containing 0.02% of sodium azide and prepared for immunofluorescence labeling.

Two common marmosets (*Callithrix jacchus*; one male and one female) weighing around 370 g were used for this study. They were caged at 27 ± 2 °C in 50 ± 10% humidity with a 12-h light–dark cycle and were fed twice a day with a standard marmoset diet supplemented with fruit, mealworm, and gum with vitamin D. Water was available *ad libitum*. Following sedation with ketamine hydrochloride (40 mg/kg, i.m.), the marmosets were deeply anesthetized with an overdose of sodium pentobarbital (50 mg/kg, i.v.) for perfusion-fixation. The marmosets were transcardially perfused with 0.1 M PBS followed by 4% paraformaldehyde in 0.1 M PB. The brain was removed from the skull, postfixed in the same fresh fixative overnight, and saturated with 30% sucrose in PB at 4 °C. Tissue blocks containing the putamen and caudate nucleus were sectioned coronally using a freezing microtome at a thickness of 25 µm. Twelve series of floating sections were collected in 0.1 M PB containing 0.02% of sodium azide and prepared for immunofluorescence labeling.

#### 2.2.2 Immunofluorescence labeling

The brain sections of mice, rats, and marmosets were incubated with a mixture of primary antibodies overnight at RT (Table 1). The primary antibodies were diluted with incubation buffer containing 10% (v/v) normal donkey serum (Merck KGaA, Darmstadt, Germany), 2% bovine serum albumin and 0.5% (v/v) Triton X-100 in 0.05 M Tris-buffered saline (TBS). After exposure to the primary antibodies, the sections were washed in TBS and incubated for 3 h at RT in the same buffer containing a mixture of secondary antibodies (Table 2). In some cases, the whole immunoreaction steps were repeated to enhance signals. After rinsing, the sections were mounted on glass slides, air dried, and cover-slipped with 50% (v/v) glycerol/TBS.

**Table 1.**
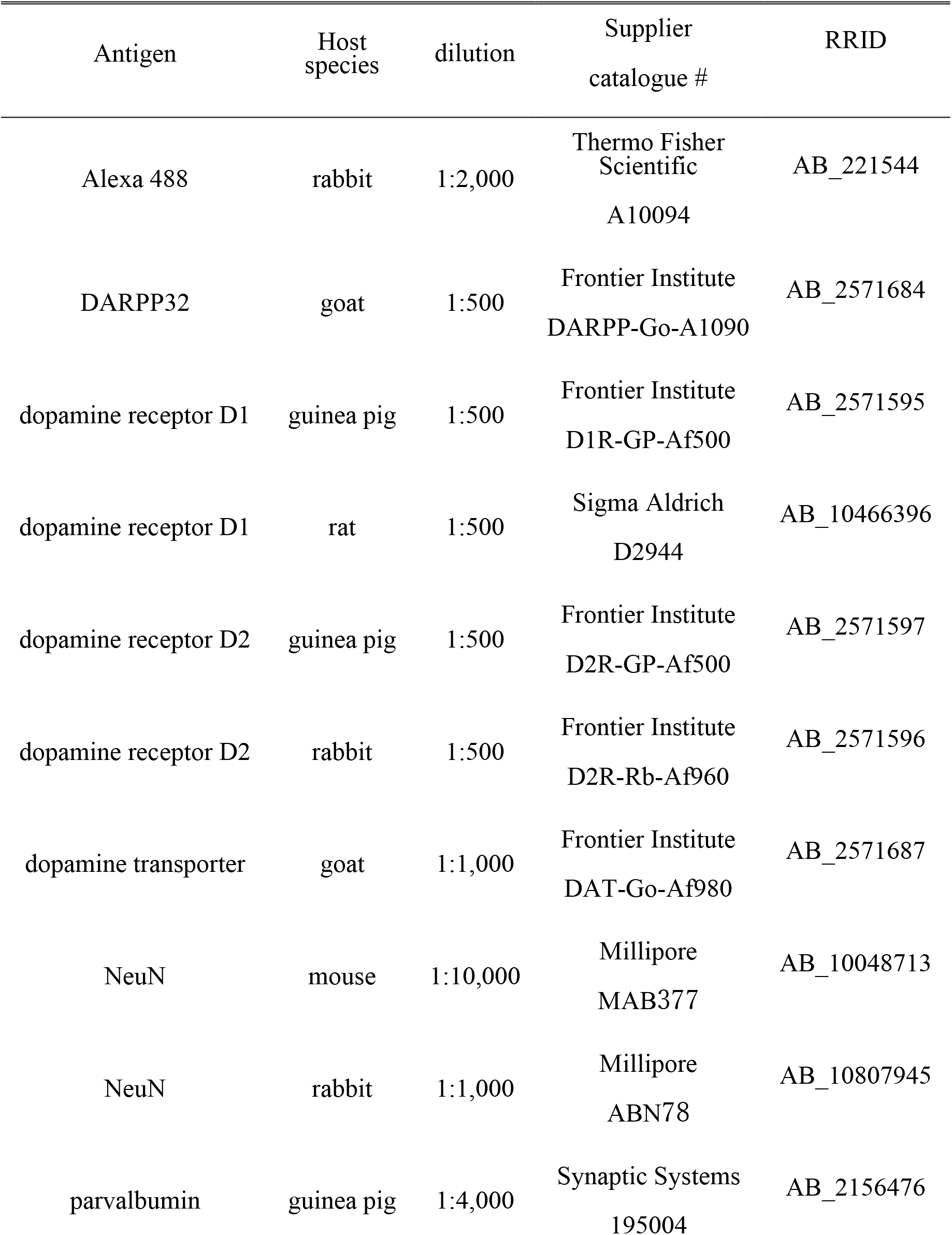
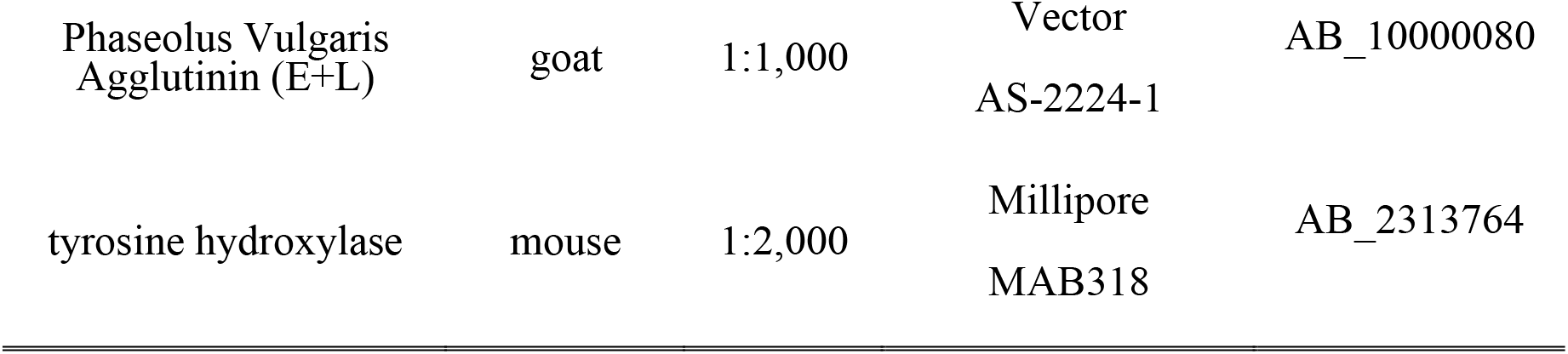
Primary antibodies used in this study

**Table 2.** Secondary antibodies and streptavidin used in this study.

| antibody | Host species | dilution | Supplier catalogue # | RRID |
| --- | --- | --- | --- | --- |
| Anti-goat Alexa Fluor 546 | donkey | 1:500 | Thermo Fisher Scientific<br>A11056 | AB_10584485 |
| Anti-goat Alexa Fluor 633 | donkey | 1:500 | Thermo Fisher Scientific<br>A21082 | AB_2535739 |
| Anti-guinea pig Alexa Fluor 488 | donkey | 1:500 | Jackson ImmunoResearch<br>706-545-148 | AB_2341098 |
| Anti-guinea pig Alexa Fluor 555 | goat | 1:500 | Thermo Fisher Scientific<br>A21435 | AB_2535856 |
| Anti-guinea pig Alexa Fluor 633 | goat | 1:500 | Thermo Fisher Scientific<br>A21105 | AB_2535757 |
| Anti-mouse Alexa Fluor 488 | donkey | 1:500 | Thermo Fisher Scientific<br>A21202 | AB_2535788 |
| Anti-mouse Alexa Fluor 546 | donkey | 1:500 | Thermo Fisher Scientific<br>A10036 | AB_2534012 |
| Anti-mouse Alexa Fluor 635 | goat | 1:500 | Thermo Fisher Scientific<br>A31575 | AB_2536185 |
| Anti-rabbit-biotin-SP | donkey | 1:100 | Jackson ImmunoResearch<br>#711-065-152 | AB_2340593 |
| Anti-rabbit Alexa Fluor<br>488 | donkey | 1:500 | Thermo Fisher<br>Scientific<br>A21206 | AB_2535792 |
| Anti-rabbit Alexa Fluor<br>546 | donkey | 1:500 | Thermo Fisher<br>Scientific<br>A10040 | AB_2534016 |
| Anti-rabbit Alexa Fluor<br>635 | goat | 1:500 | Thermo Fisher<br>Scientific<br>A31577 | AB_2536187 |
| streptavidin Alexa Fluor<br>488 | - | 1:1000 | Thermo Fisher<br>Scientific<br>S32354 | AB_2315383 |

#### 2.2.3 Tracer visualization

BDA was visualized with fluorophore-conjugated streptavidin (Thermo Fisher Scientific, Table 2; 1:1,000 for 3 h). The reaction was enhanced using the biotinylated tyramine (BT)-glucose oxidase (GO) amplification method (Furuta et al., 2009; Ge et al., 2010; Kuramoto et al., 2009). PHAL was detected using immunofluorescence.

### 2.3 *In situ* hybridization and NeuN immunolabeling

All *in situ* hybridization experiments were accomplished under a ribonuclease-free condition. Mice (C57BL/6J) for *in situ* hybridization were treated as aforementioned in section 2.2, except, picric acid was excluded from the fixative. The following hybridization procedure was performed as reported previously (Hioki et al., 2010; Ma et al., 2011). Briefly, sagittal sections from both hemispheres were cut at 20-μm thickness using a freezing microtome. Free floating sections were hybridized for 16—20 h at 60 ℃ with 1 μg/mL digoxigenin (DIG)-labeled sense or antisense riboprobes in a hybridization buffer. After washes and ribonuclease A (RNase A) treatment, the sections were incubated overnight with 1:1,000 diluted alkaline phosphatase-conjugated anti-DIG sheep antibody (11-093-274-910; Roche Diagnostics, Basel, Switzerland) and then reacted with 0.375 mg/mL nitroblue tetrazolium and 0.188 mg/mL 5-bromo-4-chloro-3-indolylphosphate (NBT/BCIP; Roche Diagnostics) for 27—42 h.

Sense probes detected no signal higher than the background. To sensitively detect the signals for Drd1a and Drd2 mRNA, we also applied BT-GO amplification method as aforementioned in section 2.2.3. Briefly, after hybridization with DIG-labeled Drd1a and Drd2 riboprobes, the sections were incubated with 1:4,000 diluted peroxidase-conjugated anti-DIG sheep antibody (11-207-733-910; Roche Diagnostics). Subsequently, the sections were reacted with a mixture containing 31 µM BT, 3 µg/mL of GO, 2 mg/mL of beta-D-glucose, and 2% bovine serum albumin in 0.1 M PB for 30 min. The sections were further incubated with 1:1,000 diluted alkaline phosphatase-conjugated streptavidin (02516-71; Nacalai Tesque) for 2 h and finally reacted with NBT/BCIP. The probes for Drd1a (target sequence position, 1116-1809 GenBank: NM_010076.3, gifted from Dr. Shinichiro Okamoto) and for Drd2 (target sequence position, 1412-2497 GenBank: X55674.1, gifted from Dr. Shinichiro Okamoto) were used.

After *in situ* hybridization, the sections were processed for NeuN immunohistochemistry for conventional visualization with bright microscopy using avidin-biotin-peroxidase complex (ABC Elite; Vector, Burlingame, CA) and diaminobenzidine. The stained sections were serially mounted onto the gelatinized glass slides, dried, washed in running water, dried again, cleared in xylene, and finally covered with mounting medium MX (Matsunami, Kishiwada, Japan) and a glass coverslip. The boundaries of the D1R- and D2R-poor zones were determined with double immunofluorescent staining for D1R and D2R using the adjacent section.

### 2.4 Electrophysiological recording and post-hoc immunofluorescence

The electrophysiological recording using *in vitro* slice preparation was conducted as previously reported (Karube et al., 2019). Briefly, a mouse (postnatal 3—4 weeks) was deeply anesthetized with isoflurane and decapitated. The brain was removed from the skull and immediately cooled for 2 min in ice-cold artificial cerebrospinal fluid (ACSF) oxidized with 95% O2/5% CO2 gas. Then, a brain block containing the striatum was resected and coronally sectioned into slices of 300 µm thickness using a vibratome (7000smz-2, Campden, Leicestershire, UK) in cold ACSF. The sections were incubated for 20 min at 32 ℃ and then over 1 h at RT for recovery. Striatal neurons and neurons in the neighboring nuclei were recorded using the whole cell patch clamp method with the aid of an EPC10 amplifier (HEKA Elektronik Dr. Schulze GmbH, Lambrecht/Pfalz, Germany). The pipette solution was composed of K-gluconate 130; KCl2; Na2ATP 3; NaGTP 0.3; MgCl2 2; Na4EGTA 0.6; HEPES 10; biocytin 20.1 (in mM). The pH was adjusted to 7.3 with KOH, and the osmolality was ∼290 mOsm. To obtain the relationship between action potential frequency and input current intensity, 1000 ms of depolarized current pulses were applied with a 50 pA-increment step. From the data, rheobase current was calculated. After electrophysiological recording, an image of the slice and electrode tip was acquired to clarify the location of the recorded neurons in the striatum. The slices were fixed overnight with a fixative composed of 4% paraformaldehyde and 0.2% picric acid dissolved in 0.1 M PB. The fixed slices were re-sectioned into 50 µm slices, and the recorded neurons filled with biocytin were visualized using fluorophore-conjugated streptavidin. D1R, D2R, and DARPP32—a marker of MSNs—were identified by immunofluorescent labeling as described above (section 2.2) to confirm the location of the recorded neurons, and to confirm if they were MSNs. The image was aligned to the corresponding image taken at recording to identify individual neurons.

### 2.5 Image acquisition and processing

The specimens were observed using the following microscopes: all-in-one fluorescence microscope (BZ-X710; Keyence, Osaka, Japan); BX53 (equipped with a DP73 CCD camera; Olympus, Tokyo, Japan) or a confocal microscope (FV1200, Olympus). For fluorescent imaging, appropriate filter sets (359–371-nm excitation and 397-nm emission for Alexa Fluor (AF) 350 or 405; 450–490-nm excitation and 514–565-nm emission for AF488; 530–585-nm excitation and 575–675-nm emission for AF594; 590–650-nm excitation and 655–675-nm emission for AF635) were applied. The images of each channel were obtained sequentially and separately to negate possible crosstalk of signals across channels. Sections processed for *in situ* hybridization were observed with bright field microscopy.

To quantify and compare immunofluorescent signals for D1R, D2R, and tyrosine hydroxylase (TH) across the striatal regions and among individual mice, first D1R- and D2R-poor regions were approximately determined using immunofluorescent images of a sagittal section. A line was drawn to connect the ventral border of the internal capsule (ic) and the mid point of the rostral edge of D1R- poor zone, then the line was extended toward the cerebral cortex in a linear fashion (Figure 2B).

Finally, a pixel intensity profile along the line was obtained. The precise borders between the rostral striatum and D1R-poor zone, D1R-poor zone and D2R-poor zone, or D2R-poor zone and ic, were confirmed using the derivative of each line plot as the point of maximum slope. Finally, regions of interest (ROIs, 200 × 200 µm^2^ unless otherwise noted) were placed in each region along with the line. To measure pixel intensity of the ROIs, small areas containing nerve bundles were masked.

For quantification of anterogradely labeled axon distribution (Figures 6, 7), the acquired confocal images (1 µm of Z-spacing) were analyzed using Fiji—an image processing package (Schindelin et al., 2012). The sections were processed for triple fluorescent labeling for BDA- or PHAL-loaded axons, TH, and NeuN in every 120-µm coronal sections. Using fluorescence of TH, NeuN, and autofluorescence of thick axon bundles (eg. cerebral peduncle), the contours of GP and SNpl were delineated. The image containing axons was processed with “subtract background” function of Fiji to reduce background fluorescence, and then Z projection of the image was obtained. Next, the image was thresholded and binarized. The number of pixels containing axons were counted using the binarized image. In individual mice, the labeled axons were contained in 2– 4 GP sections, and in 3– 5 SNpl sections. Since the number of the axon pixels varied among sections, especially for GP, the maximum number of pixels per section was compared between axons in GP and axons in SNpl.

### 2.6 Statistical comparison

All averaged values are represented as mean ± standard deviation. The quantitative values among groups (> 2) were compared using one-way ANOVA followed by the post-hoc Tukey’s test with the aid of software: Microsoft Excel, R (language and environment for statistical computing and graphics similar to S, http://cran.r-project.org/), and MATLAB (MathWorks, Natick, MA). Comparisons between two data groups were conducted using t-test. All p-values are presented; p-values less than 0.05 were considered to be statistically significant.

## 3 Results

### 3.1 Uneven distribution of immunoreactivity of D1R and D2R in the mouse caudal striatum

In marked contrast to the dorsal, rostral, or medial striatum, D1R-immunoreactivity was weak from 1.1 mm to 2.0 mm posterior to the bregma (AP −1.1 – −2.0) and from 2.7 mm to 3.3 mm lateral to the midline (L 2.8 – 3.3), whereas D2R-immunoreactivity was weak in the range of AP −1.2 – −1.8 and L2.7 – 3.2 in mice. Representative immunofluorescent images for D1R, D2R, and TH on sagittal and coronal planes of the striatum are shown in Figure 1. Uneven distributions of D1R, D2R, and TH were obviously observed in the caudal (AP −1.3, −1.6, and −1.9 in coronal plane in Figure 1A) and lateral (L 2.9, 3.0, 3.2, and 3.3 in sagittal plane in Figure 1B) part of the striatum. The zone with weak D1R and that with weak D2R separately existed (arrowhead in Figure 2A; cf. Gangarossa et al., 2013b; Miyamoto et al., 2019). Hereafter, they are referred to as D1R-poor zone and D2R-poor zone, respectively. In these zones, immunofluorescence for TH was also weak (Figures 1, 2A; Miyamoto et al., 2019). A representative sagittal section (L3.0) and the definition of the striatal area used in this study are shown in Figure 2A and 2B, respectively. This characteristic spatial expression of D1R, D2R, and TH was always observed in all samples used for this study (N = 45 mice). To quantify these changes of fluorescence, fluorescence intensity line profiles were obtained for D1R, D2R, and TH as shown in Figure 2B (N = 3 mice; one section/mouse; see Materials and Methods 2.5). The fluorescence intensity was compared in ROIs located at the rostral striatum (rStr), D1R-poor zone, and D2R-poor zone, and then normalized by the values of rStr (Figure 2C). Normalized D1R pixel intensity was significantly different among the zones as analyzed by one-way ANOVA (F = 104.25, *p* = 2.2 × 10^−5^). D1R expression was lower in D1R-poor zone (0.53 ± 0.02) than rStr (*p =* 0.00002) or D2R-poor zone (0.92 ± 0.06; *p =* 0.00008) with post-hoc Tukey’s test. D2R was also expressed differentially among zones (F = 131.44, *p* = 1.1 × 10^−5^ by one-way ANOVA). In contrast to D1R, D2R expression was significantly lower in D2R-poor zone (0.33 ± 0.08) than rStr (*p =* 0.00003 with post-hoc Tukey’s test) or D1R-poor zone (1.09 ± 0.03; *p =* 0.00001). In addition, TH expression differed among zones (F = 336.89, *p* = 6.9 × 10^−7^ by one-way ANOVA). The pixel intensity was lower in D1R-poor zone and D2R-poor zone (0.41 ± 0.04 and 0.44 ± 0.03, respectively) than rStr (*p =* 0.000001 for D1R-poor zone and *p =* 0.000002 for D2R-poor zone). On the other hand, DARPP32, a well-known marker of MSNs, was expressed in the whole area of the striatum including the poor zones (Figure 2D). At higher magnifications, immunofluorescence of DARPP32 in cell bodies was almost similar throughout the striatum (Figure 2E), although it was apparently weak in cStr at low magnification (Figure 2D). When combined with NeuN-immunostaining, the number of DARPP32-expressing neurons was counted in ROIs placed in four striatal subregions (ROI size, 318 × 318 µm^2^; Figure 2D, 2E; N = 3 mice, one section/mouse). The proportion of DARPP32-expressing cells to the number of NeuN-positive cells was 94.3 ± 1.3% in rStr (N = 437/464 NeuN-expressing cells), 93.5 ± 0.9% in para-poor zone (ppz, N = 392/419), 92.6 ± 0.5% in D1R-poor zone (N = 465/502), and 92.1 ± 0.5% in D2R-poor zone (N = 396/430) (Figure 2F). The proportion was not significantly different among rStr, para-poor zone, and two poor zones (F = 2.95, *p* = 0.15, analyzed by one way ANOVA).

**Figure 1.**
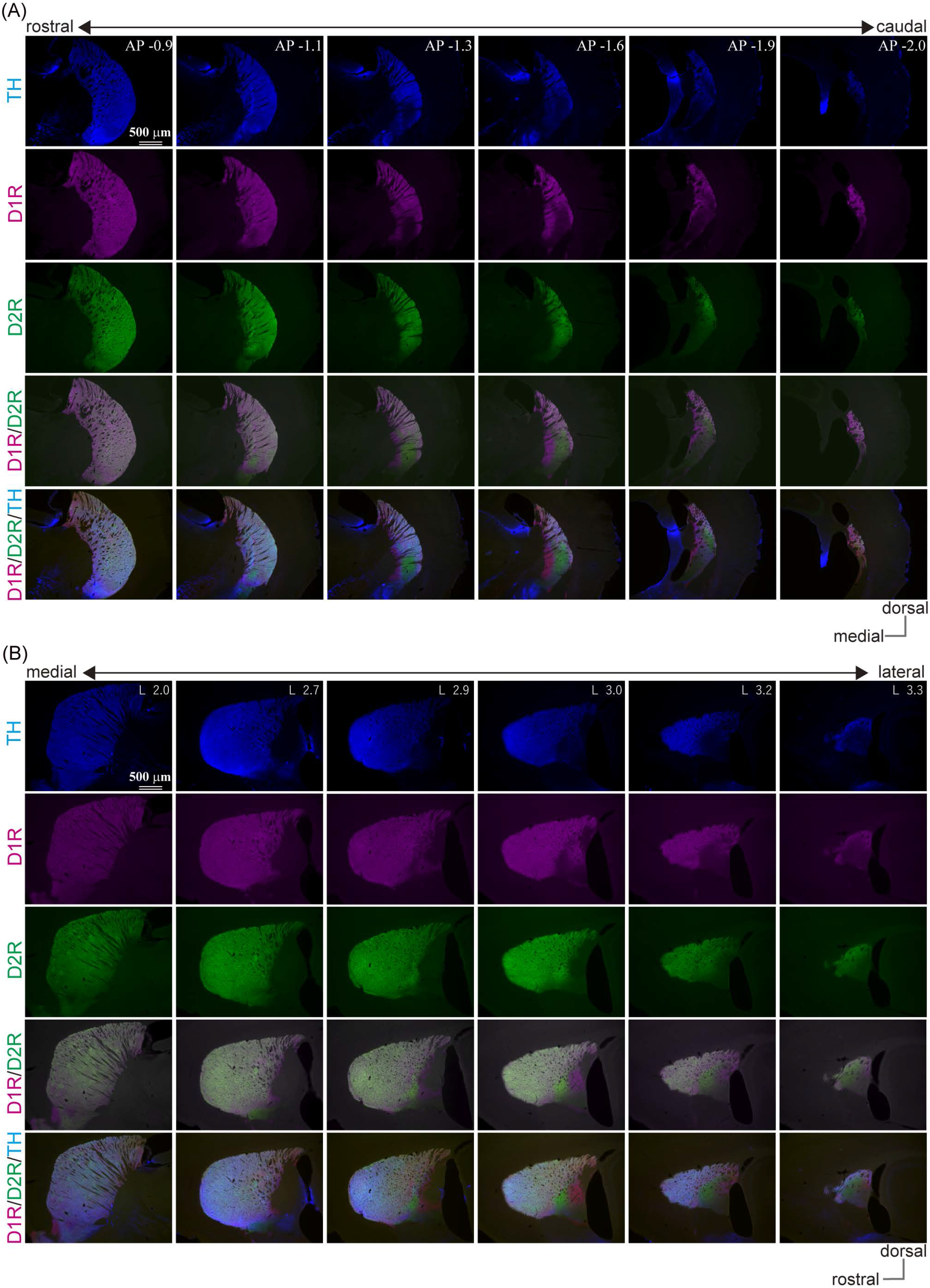
Uneven distribution of immunofluorescence for dopamine receptor D1 (D1R), dopamine receptor D2 (D2R), and tyrosine hydroxylase (TH) in the mouse caudal striatum. Images of immunofluorescent labeling against TH (blue), D1R (magenta), and D2R (green) show explicit location of the D1R/D2R-poor zones in the mouse striatum. (A) Coronal plane. The number seen after the abbreviation “AP” at the right upper corner of the top-row images represents “distance from the bregma” in mm. (B) Sagittal plane. The number seen after the abbreviation “L” at the right upper corner of the top-row images represents “lateral distance from the midline” in mm.

**Figure 2.**
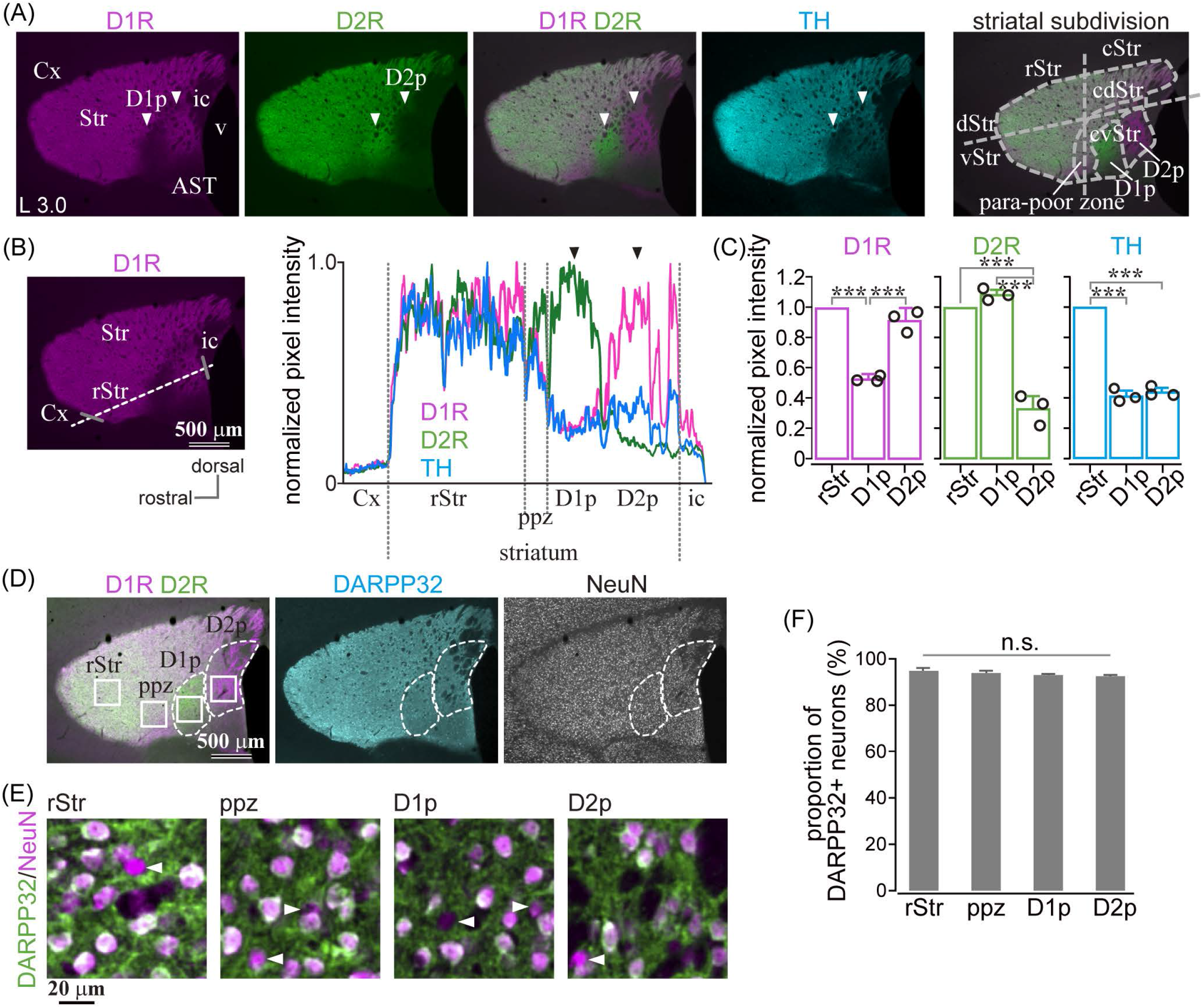
Quantification of dopamine receptor D1 (D1R), dopamine receptor D2 (D2R), and tyrosine hydroxylase (TH) expression in the caudal striatum. (A) Images of immunofluorescent labeling for D1R (magenta), D2R (green), and TH (cyan) of the mouse striatum. Arrowheads indicate the regions with very faint signal as either D1R or D2R. For this paper, the lateral striatum is subdivided into the rostral (rStr) and caudal striatum (cStr); the dorsal (dStr) and ventral striatum (vStr); cStr is further divided into the caudo-dorsal (cdStr) and caudoventral striatum (cvStr), as shown in the rightmost panel. D1R- and D2R-poor zones (abbreviated as D1p and D2p, respectively) were situated in cvStr. The region surrounding the poor zones is named here as para-poor zone in which nearly uniform expression of D1R and D2R is seen. (B) Quantification of pixel intensity along a dotted white line. Gray bars represent the rostral and caudal edges of the striatum. The right panel shows normalized pixel intensity in a section of a mouse (3.0 mm lateral from the midline). (C) Average pixel intensity (N = 3 mice). Circles indicate individual data. (D) Quadruple immunostaining against D1R, D2R, NeuN, and DARPP32. DARPP32 was expressed in the whole striatum (cyan). (E) Magnified images of NeuN (magenta) and DARPP32 (green) immunostaining. To clearly represent overlay of NeuN and DARPP32, the pseudocolors were different from those shown in D. Arrowheads indicate neurons without DARPP32 expression. (F) Proportion of DARPP32 expressing neurons. AST, amygdala striatal transition area; Cx, cerebral cortex; ic, internal capsule; v, ventricle. \*\*\**p* < 0.001.

Dopamine transporter (DAT) is purely expressed in dopaminergic nervous system, while TH expresses in catecholaminergic axons as well as dopaminergic axons. Thus, we examined whether DAT expression showed similar spatial pattern to TH. As shown in Supplementary Figure 1, distributions of DAT and TH highly overlapped in the whole striatum, except for subcallosal zones where TH expression was relatively low. The line profile for DAT immunofluorescence correlated highly with that for TH (Supplementary Figure 1B, Pearson’s correlation coefficient was 0.953 ± 0.012, *p* < 2.2 × 10^−6^; N = 3 mice, one section/mouse). DAT immunofluorescence was significantly lower in the caudo- ventral striatum (cvStr) than rStr (Supplementary Figure 1C; *p* = 0.03125; N = 2 ROIs for each rStr and cvStr in each section; N = 3 mice; one section/mouse). Normalized intensity in cvStr was 0.475 ± 0.111 for DAT and 0.457 ± 0.114 for TH.

### 3.2 Uneven distribution of mRNA of D1R and D2R in caudal striatal neurons in mice

Since D1R and D2R are also expressed in presynaptic terminals, it is not clear whether the above investigation reflects differential cellular compositions of specific area of the cStr. Even if the fluorescence mainly came from cell bodies, different fluorescence intensities could reflect total neuronal density among striatal regions. To directly solve the question, we investigated mRNA expression of Drd1a and Drd2 genes by *in situ* hybridization. A section containing the poor zones was used for *in situ* hybridization with NeuN immunostaining. To confirm the poor zone identity, the adjacent section was used for double immunofluorescent staining of D1R and D2R (Figure 3A, 3B, 3D, 3E). Drd1a (Figure 3A, 3B) and Drd2 (Figure 3D, 3E) were detected (dark blue) in cell bodies of striatal neurons expressing NeuN (brown). To compare zonal differences in the number and proportion of D1R- or D2R-neurons, we counted them in four areas: rStr, para-poor zone, D1R-poor zone, and D2R-poor zone (see Figure 2A for subdivision of the striatum used in this study). The number of NeuN immunopositive cells did not significantly differ among four ROIs (N = 3 mice; one section/mouse; N = 5,689 neurons in total; F = 1.26, *p* = 0.14 by one-way ANOVA; Figure 3G), suggesting that the cell density of the striatum, most of which is probably regarded as the MSNs density (Figure 2F), seems to be uniform even in the two poor-zones. The number of Drd1a or Drd2 expressing neurons were counted in one representative section for each mouse (N = 3 mice; one section/mouse; N = 2,615 neurons for Drd1a and N = 3,074 neurons for Drd2). The proportion of Drd1a-expressing neurons to total neurons was 12.23 ± 1.43% (N = 73/593 neurons as sum of three mice) in D1R-poor zone, 82.06 ± 3.08% (N = 528/645) in D2R-poor zone, 50.44 ± 0.07% (N = 354/702) in rStr, and 46.49 ± 1.34% (N = 315/675) in para-poor zone, which were significantly different among these regions (F = 579.5; *p* = 0.0000507 using one-way ANOVA). Post-hoc Tukey’s test revealed that the proportion of Drd1a-expressing neurons was significantly lower in D1R-poor zone (*p* = 0.0000001 *vs*. rStr; *p* = 0.0000001 *vs.* para- poor zone; *p* = 0.0000001 *vs.* D2R-poor zone), and significantly higher in D2R-poor zone than the other three regions (*p* = 0.0000001 *vs.* rStr; *p* = 0.0000001 vs para-poor zone) (Figure 3C). No significant difference was observed for rStr *vs*. para-poor zone (*p* = 0.13). Contrastingly, the proportion of Drd2-expressing neurons to total neurons was 78.68 ± 1.73% (N = 539/686) in D1R-poor zone, 3.58 ± 1.19% (N = 28/786) in D2R-poor zone, 46.78 ± 4.23% (N = 366/781) in rStr, and 46.73 ± 4.13% (N = 382/821) in para-poor zone. The proportion was significantly different among these four regions (F = 998.48; *p* = 0.00000276 using one-way ANOVA). D2R-poor zone contained a significantly lower proportion of D2R-expressing neurons than the other regions (*p* = 0.0000007 *vs.* rStr; *p* = 0.0000007 *vs.* para-poor zone; *p* = 0.0000001 *vs.* D1R-poor zone). D1R-poor zone contained a significantly higher number of D2R-expressing neurons than others (*p* = 0.0000076 *vs.* rStr; *p* = 0.0000075 *vs.* para-poor zone) (Figure 3F). Again, no significant difference was observed between rStr and para-poor zone (*p* = 0.99). Since we could not visualize both Drd1a and Drd2 simultaneously, their composition in each area was elucidated as the sum of individual data in Figure 3H. These results clearly indicate the distinct cell type composition of both D1R- and D2R-poor zones, in which either D2R or D1R expressing neurons occupied approximately 80% of total neurons. In D2R-poor zone, proportion of neurons expressing neither D1R nor D2R was slightly larger than other zones. We also compared interneuron density for neurons expressing parvalbumin, calretinin, or choline acetyl transferase. As Miyamoto et al. (2019) reported, subtle differences were observed. However, this information is still insufficient to explain the proportion of neurons that lack D1R/D2R expression in the D2R-poor zone.

**Figure 3.**
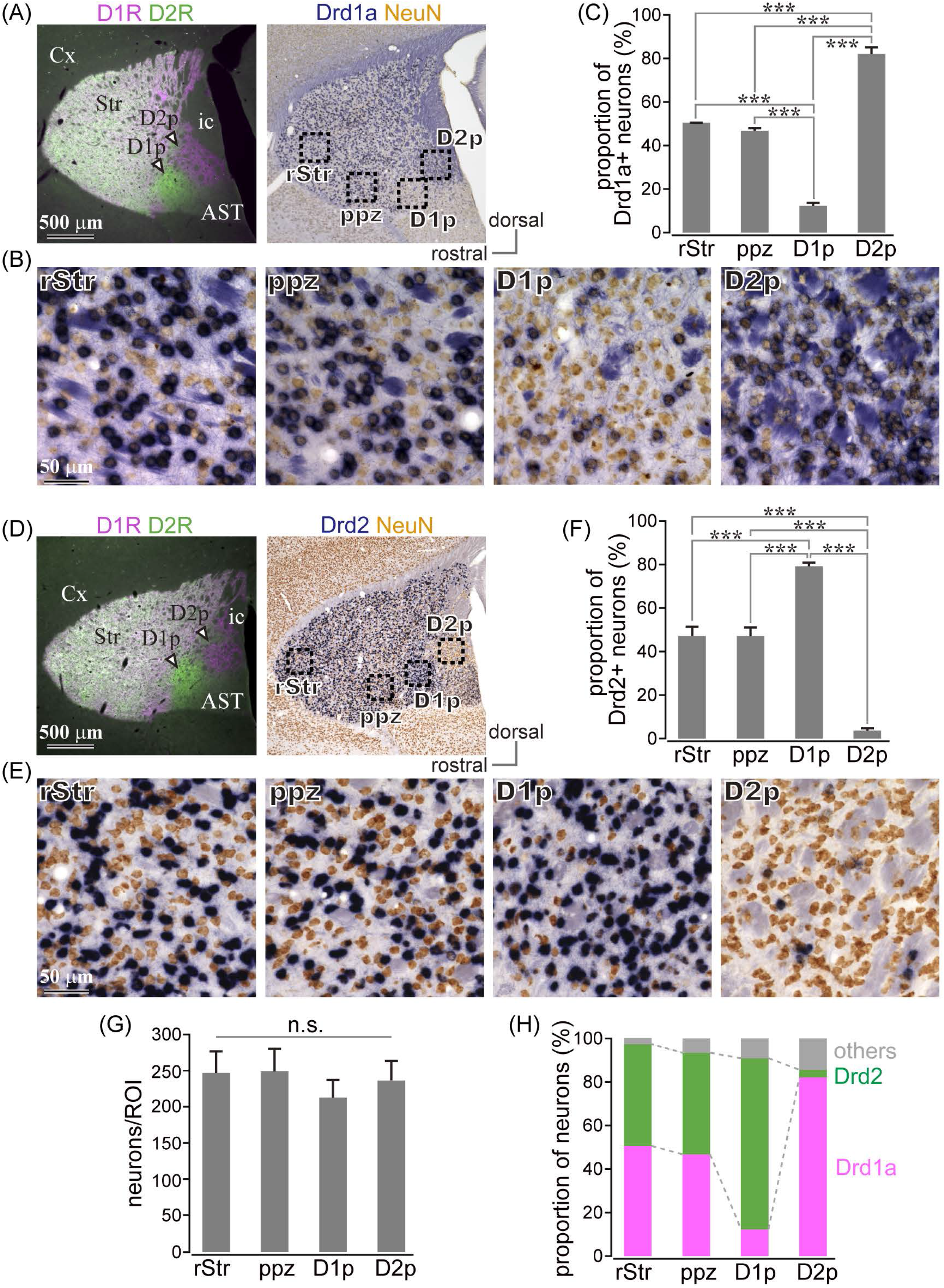
Uneven distribution of dopamine receptor D1 (D1R) or dopamine receptor D2 (D2R) mRNA expressing neurons in the caudal striatum. (A), (D) Left, Images of immunofluorescent labeling for D1R (magenta) and D2R (green). Right, Drd1a- (A) or Drd2- (D) expressing neurons (blue) with immunostaining against NeuN (brown). (B), (E) Magnified images of rectangle areas 1–4 in A or D. (C), (F) Proportion of Drd1a- or Drd2- expressing neurons in rostral (rStr), para-poor zone (ppz), D1R-poor zone (D1p) or D2R-poor zone (D2p). (N = 3 mice for each) (G) Mean NeuN+ cell number in each ROI. (H) Elucidated cell composition in each striatal area. AST, amygdala striatal transition area; Cx, cerebral cortex; ic, internal capsule; v, ventricle. \*\**p* < 0.01.

### 3.3 MSNs in D1R- or D2R-poor zones possessed similar membrane properties to those in the dorsal striatum

Since the poor zones are located close to the boundary between the striatum and the amygdala-striatal transition area (AST) or GP, we wondered whether these dopamine-receptor expressing neurons are MSNs possessing similar properties to other parts of the striatum. Whole cell patch clamp recordings were accomplished from medium sized neurons in those area (N = 8 mice), and the recorded neurons were filled with biocytin. The recorded slices were examined for post-hoc immunofluorescence against D1R, D2R, and DARPP32 (Figure 4A, 4B). As a result, we obtained medium sized neurons recorded in the cdStr (N = 6) and D1R- (N = 5) or D2R-poor zone (N = 3) (Figure 4D). They represented similar membrane properties, such as deep resting membrane potentials, narrow action potentials, and low input resistances (*p* > 0.05 analyzed by one-way-ANOVA; Figure 4C, 4E, 4F, Table 3). F-I curves, relationship between action potential frequency and input current intensity, were plotted in Figure 4E.

**Figure 4.**
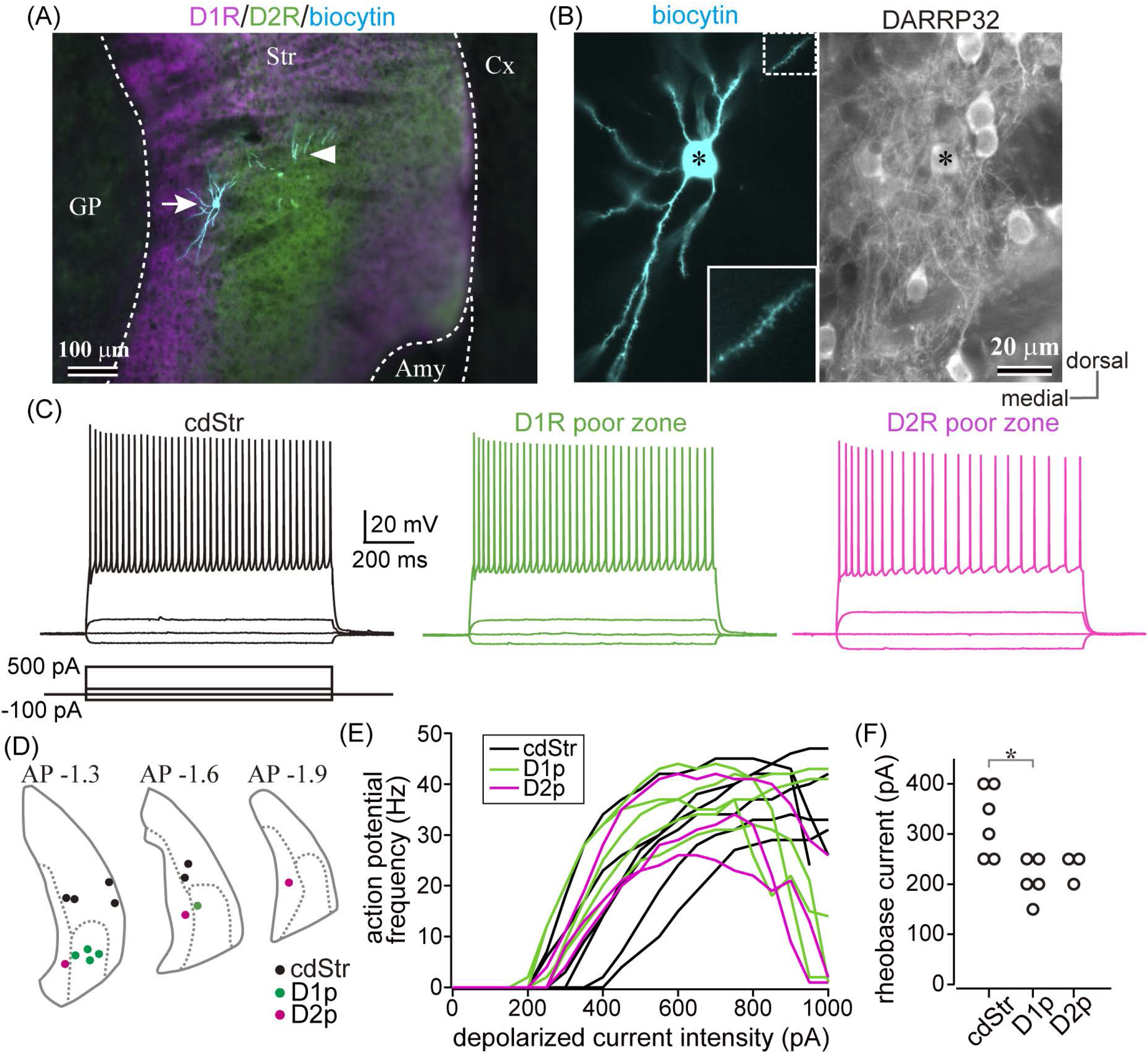
Whole cell recording from the caudal striatum. (A), An example of the whole cell recorded neurons. The location of the recorded neurons was confirmed using post-hoc immunofluorescence for dopamine receptor D1 (D1R) (magenta) and dopamine receptor D2 (D2R) (green). Recorded neurons are shown in cyan. The left neuron (arrow) is situated in D2R-poor zone (D2p), whereas the dendrites of the right neuron (arrowhead) are in D1R- poor zone (D1p). The cell body of the right neuron was not confined in this section. (B) A magnified image of the left neuron in A. Left, note many spines protruded from the dendrites. Inset shows further magnified view of the dotted rectangle area. Right, the neuron (asterisk) expressed DARPP32, a marker of MSNs. (C) Representative traces of membrane voltage responses to depolarized (100 and 500 pA) and hyperpolarized (-100 pA) current pulses. The traces were recorded from the neurons in the caudo- dorsal striatum (cdStr, left), D1R-poor zone (middle), and D2R-poor zone (right) (see also Table 3). (D) The locations of all recorded neurons were plotted in three coronal planes (1.3, 1.6, and 1.9 mm posterior to the bregma) of the caudal striatum. (E) Relationship between input current intensity and action potential frequencies. The initial slope was not significantly different among striatal regions. (F) Intensity of rheobase current (a 1000 ms-pulse) to induce an action potential. \**p* < 0.05.

**Table 3.**
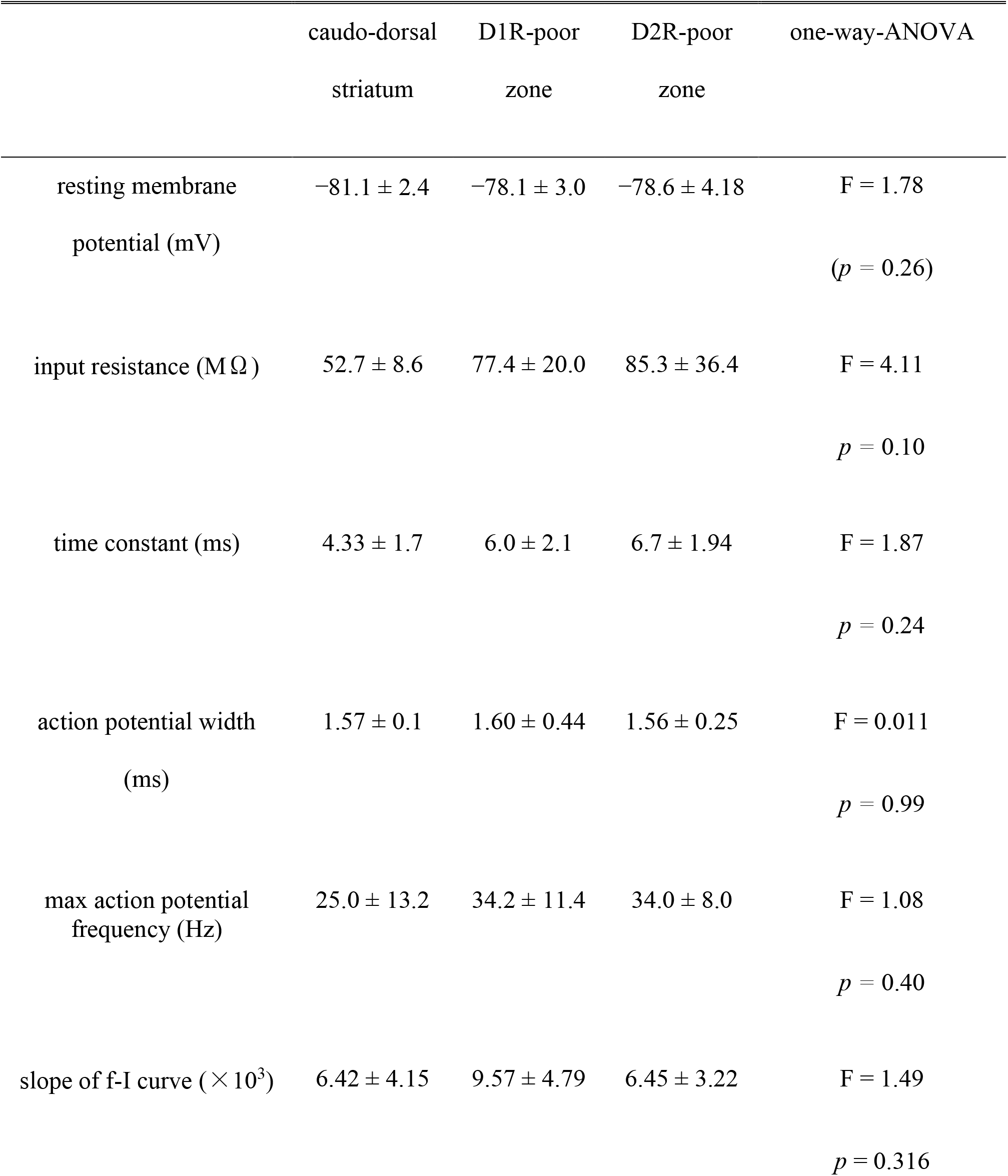
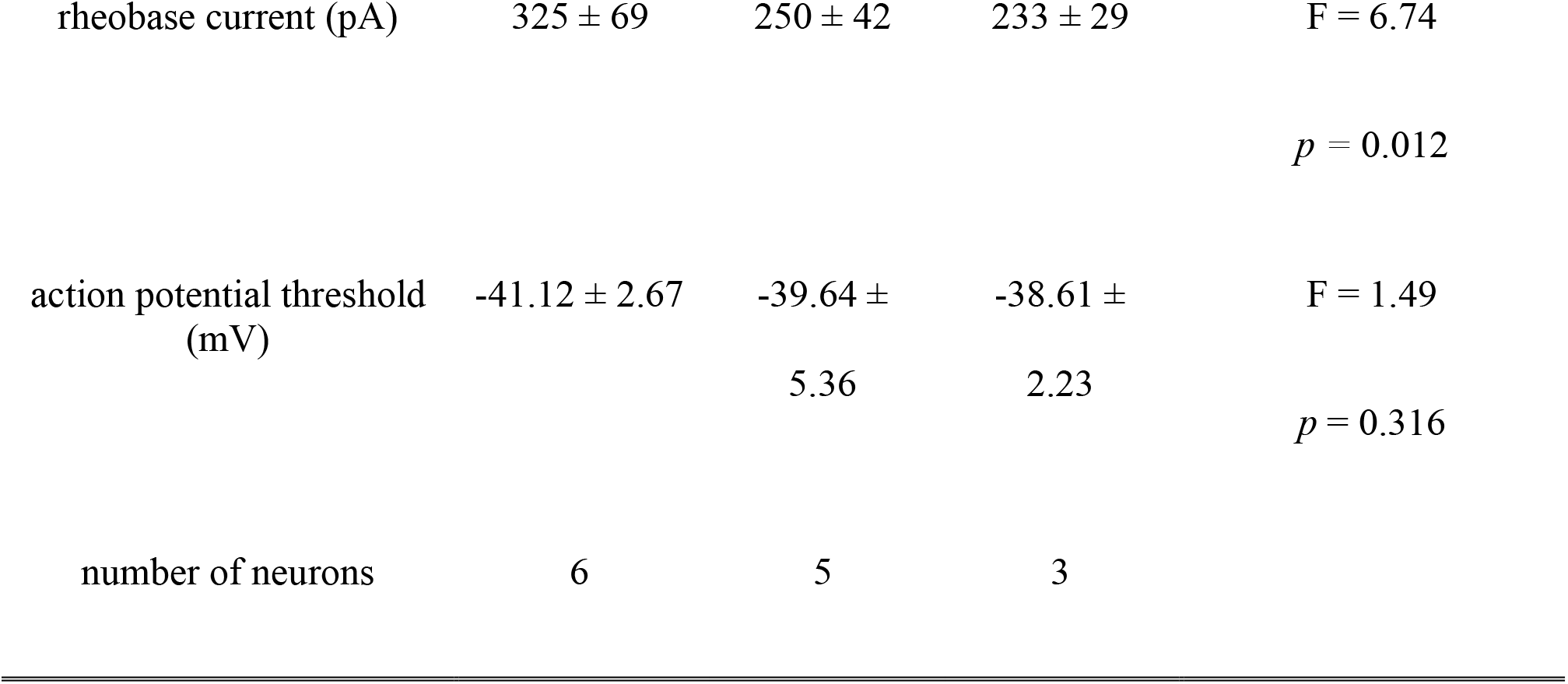
Electrophysiological properties of medium sized neurons in the dorsal striatum, D1R or D2R poor zones.

Once action potentials were elicited, the frequency increased linearly during the following 200—300 pA increments of input current. This part of the F-I curve could be linearly fitted well for all neurons (*r*^2^ = 0.971 ± 0.024). The slopes of the fitted lines were not significantly different among the recorded neurons in the three zones (Table 3, F = 1.49, *p* = 0.316 analyzed by one-way ANOVA). A significant difference was observed for rheobase current in the cdStr, D1R- and D2R-poor zones (F = 6.74, *p* = 0.012 by one-way ANOVA; Figure 4F). Post-hoc Tukey’s test indicated that rheobase current of the neurons were lower in D1R-poor zone than in cdStr (*p* = 0.013). The threshold of an action potential (elicited by a brief pulse of 10 ms) was not significantly different among three zones (F = 0.49, *p* = 0.627 by one-way ANOVA). In addition, all neurons which could be examined were positive for immunofluorescence against DARPP32 (N = 3/3 neurons in the cdStr, N = 5/5 in D1R-poor zone, and N = 3/3 in D2R-poor zone). The cell bodies of the remaining 3/6 cdStr recorded neurons were not recovered; therefore they could not be examined for DARPP32 expression. In some neurons, their dendrites were well visualized, and they were spiny (Figure 4B inset). Thus, both poor zones are composed of MSNs, similar to the cdStr.

### 3.4 Retrogradely labeled direct pathway neurons were abundantly distributed in D2R-poor zone in the mouse caudal striatum

To explore whether D1R-expressing neurons in the poor zones project to the output nuclei of the basal ganglia, i.e., the direct pathway neurons, retrograde neural tracing was conducted. To compare the proportion of retrogradely labeled neurons among striatal regions including rStr and cStr, the tracer injection should have enough volume to efficiently label the neurons regardless of their topographic projections. Therefore, a large volume of CTB555 or CTB488 was injected in wide brain regions located along the striatonigral pathway. It is crucial that the injection must be located posterior to GP, to avoid labeling of the indirect pathway neurons. As shown in Figure 5A, the tracer injection extended to the SNpr, SNpc, and EP, but not to GP. As virtually all direct pathway neurons possess axon collaterals in GP in rodents (Fujiyama et al., 2011; Kawaguchi et al., 1990; Wu et al., 2000), we did not attempt to visualize the indirect pathway neurons in the same way.

**Figure 5.**
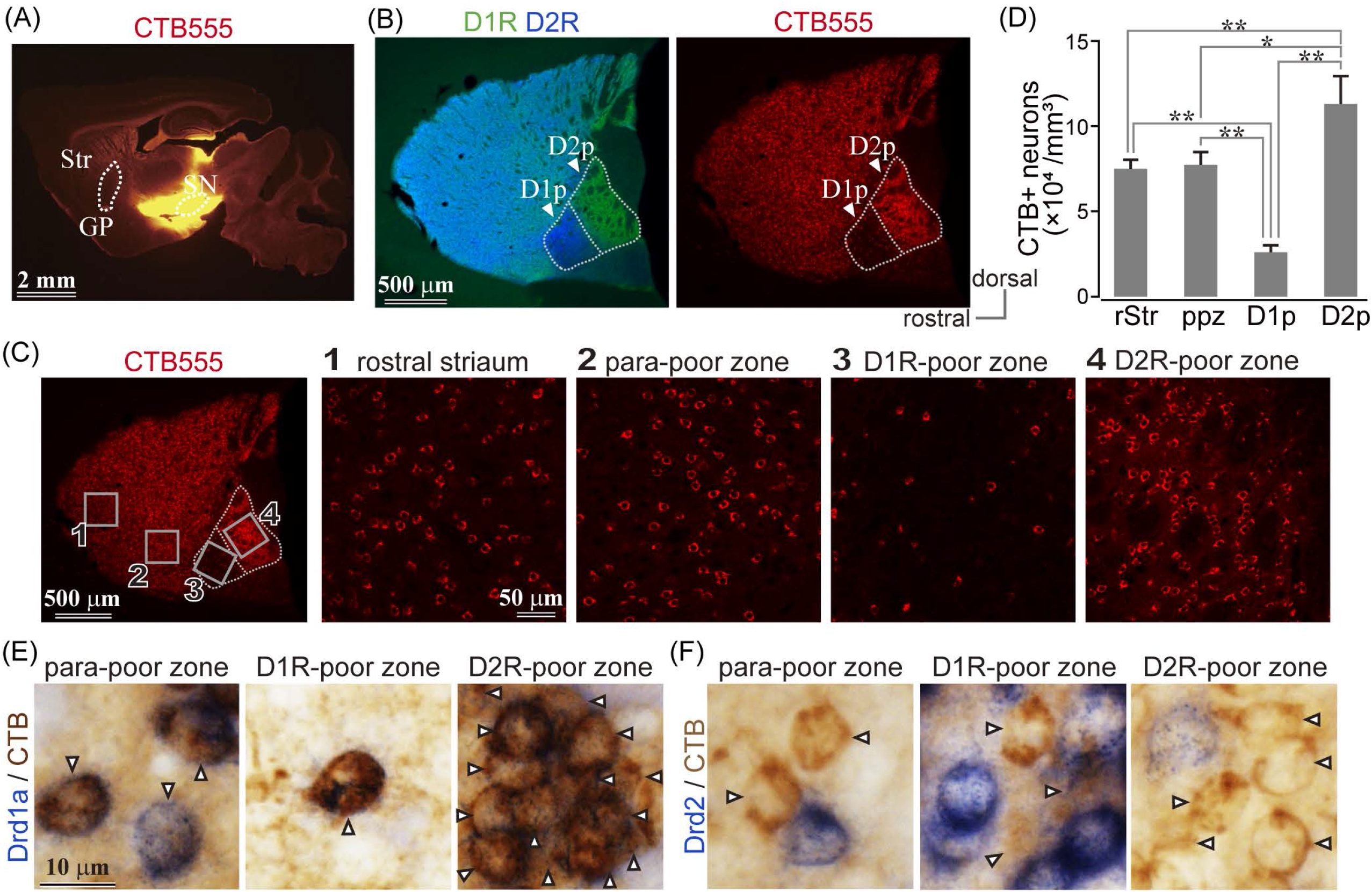
Uneven distribution of direct pathway neurons in the caudal striatum. (A) Retrograde labeling of the direct pathway neurons using bulk injection of CTB 555 in output nuclei of the basal ganglia. Note the bulk injection spread broadly; however, globus pallidus (GP), the target of the indirect pathway, is not invaded. (B) Retrogradely labeled striatal direct pathway neurons are distributed throughout the striatum, except the D1R-poor zone (D1p). (C) Magnified images of retrogradely labeled neurons in the rostral striatum, caudal striatum, D1R-poor zone, and D2R-poor zone. Their locations were indicated in the leftmost panel as rectangles 1–4. (D) Density of retrogradely-labeled neurons in each subregion. D1R-poor zone contained significantly small number of labeled neurons than the other subregions, whereas larger number of labeled neurons existed in D2R- poor zone. * *p* < 0.05; \*\**p*< 0.01. (E) Detection of Drd1a or Drd2 mRNA expression using *in situ* hybridization (blue) combined with immunohistochemistry against CTB (brown; arrowheads). Note CTB-labeled neurons expressed Drd1a (E) but not Drd2 (F).

The retrogradely labeled neurons were abundantly distributed in the whole striatum with the exception of fewer neurons in D1R-poor zone (Figure 5B). For quantification, an ROI (318 × 318 µm^2^) was sampled from each striatal region: rStr (Figure 5C1), para-poor zone (Figure 5C2), D1R-poor zone (Figure 5C3), and D2R-poor zone (Figure 5C4) (N = 3 mice; one section/mouse). In total, 2,252 retrogradely labeled neurons, which can be considered as the direct pathway neurons, were counted stereologically. The density of labeled neurons in the four striatal regions was significantly different (F = 62.392; *p* = 0.00068 using one-way ANOVA). The density was lower in D1R-poor zone (2.57 ± 0.42 × 10^4^ neurons/mm^3^) than rStr (7.52 ± 0.50 × 10^4^ neurons/mm^3^, *p* = 0.0018 using post-hoc Tukey’s test), para-poor zone (8.30 ± 1.12 × 10^4^ neurons/mm^3^, *p* = 0.00068), or D2R-poor zone (11.32 ± 1.63 × 10^4^ neurons/mm^3^, *p* = 0.00003) (Figure 5D). In contrast, the density of labeled neurons in D2R-poor zone was significantly higher than rStr (*p* = 0.009), D1R-poor zone, and para-poor zone (*p* = 0.03) (Figure 5D). These results were consistent with mRNA expression in the poor zone neurons.

In addition, to confirm that the retrogradely labeled neurons expressed D1R, we also combined *in situ* hybridization with retrograde tracing study for bright field microscopy using immunoreaction against CTB (N = 1 mouse, two sections). As shown in Figure 5E and 5F, CTB labeled neurons (brown), regarded as direct pathway neurons, expressed Drd1a (99.3%, N = 863/869) but not Drd2 (1.1%, N = 9/843) mRNA (blue).

### 3.5 Axonal projections from cdStr, D1R- or D2R-poor zones in mouse

So far, we have shown that D1R-poor zone is composed of 79% of indirect pathway neurons expressing D2R, whereas D2R-poor zone is composed of 82% of direct pathway neurons expressing D1R. A crucial question to be uncovered is whether outputs of D1R- and D2R-poor zones are similar to those of MSNs located in non-poor zones. To visualize their axonal projections, small volume of anterograde tracer (BDA, 10 kD or PHAL) was injected into the cStr, including D1R- and D2R-poor zones. Due to the small extent of the poor zones, the deposit of BDA should be extremely small to avoid spreading the tracer to neighboring striatal regions (Figures 6A, 7A–7D). Using immunofluorescence for D1R and D2R, locations of injection sites were examined. Consequently, we obtained four cases of injection to D1R-poor zone (Figures 6A1, 7B) and one case of injection to D2R-poor zone (Figure 7D). In one remaining case, injection was centered at the border between D1R- and D2R-poor zones (Figure 7C). Therefore, the projection profiles from the poor zones reported here were mainly derived from D1R- poor zone. For cdStr, the size of injection was larger compared to the poor zones as observed on fluorescent labeling (Figures 6A2, 7A; N = 3 mice). For all injections into cStr (cdStr and the poor zones), the labeled axons were mainly found in GP (Figure 6B, Supplementary Figure 2) and SNpl (Figures 6C, 7A–7D, Supplementary Figure 3), and probably, they were targets of the indirect and direct pathway MSNs, respectively. Notably, GP and SNpl were prominent targets of the cStr projections, whereas only a few collaterals were observed in EP, SNpr, and SNpc (Supplementary Figures 2, 3). We counted the number of pixels containing the labeled axons in GP and SNpl using the binarized image in every 120-µm of coronal sections (Figures 6B, 6C, 7A–7D). The anteroposterior extent of axons in GP were restricted in the 2–3 sections containing the caudal GP, whereas axons in SNpl was observed in 3–5 sections. As the axons were much more numerous in only one section for GP, we used the maximum number of axon pixels among multiple sections as an index to represent strength of projection for both GP and SNpl. The indirect pathway bias, which is the ratio of the maximum number of axon pixels in GP to that in the sum of axons in GP and SNpl, was calculated (Figure 6D). Injection into D1R-poor zone resulted in a larger indirect pathway bias than that caused by injection into cdStr (*p* = 0.031 by t-test). Injections into D2R-poor zone and into the border between D1R- and D2R-poor zone were excluded from the comparison because only one case for each was obtained. It is suggested that the indirect pathway projections to GP were relatively intensive from D1R-poor zone than from cdStr. In comparison, cdStr axons tended to be distributed in the ventral part of SNpl (Figure 7A), whereas the poor zones tended to project to the dorsal part of SNpl (Figure 7B– 7D), as reported previously for cdStr and cvStr (Foster et al., 2021; Hintiryan et al., 2016; Hunnicutt et al., 2016). To quantify and compare axon distribution in SNpl along with dorso-ventral axis, SNpl in each section was divided into 10 equally bins, then the proportion of axon pixels in each bin to total axon pixels in SNpl of the section was calculated and plotted in Figure 7E [reddish lines for D1R-poor zone projections (N = 4 mice, 16 sections) and bluish lines for cdStr projections (N = 3 mice, 11 sections)]. In the first and second bins, which are located at the most dorsal part of SNpl, the proportion was significantly larger for D1R-poor zone projections than cdStr projections (*p* = 0.0025 for the first bin by t-test; *p* = 0.0011 for the second bin). In the middle of SNpl, the proportion of cdStr projection was larger than that of D1R-poor zone projection (*p* = 0.0234). For projection to GP, the axons of both poor zones and cdStr were distributed in the caudal GP, not in the rostral GP (Supplementary Figure 2).

**Figure 6.**
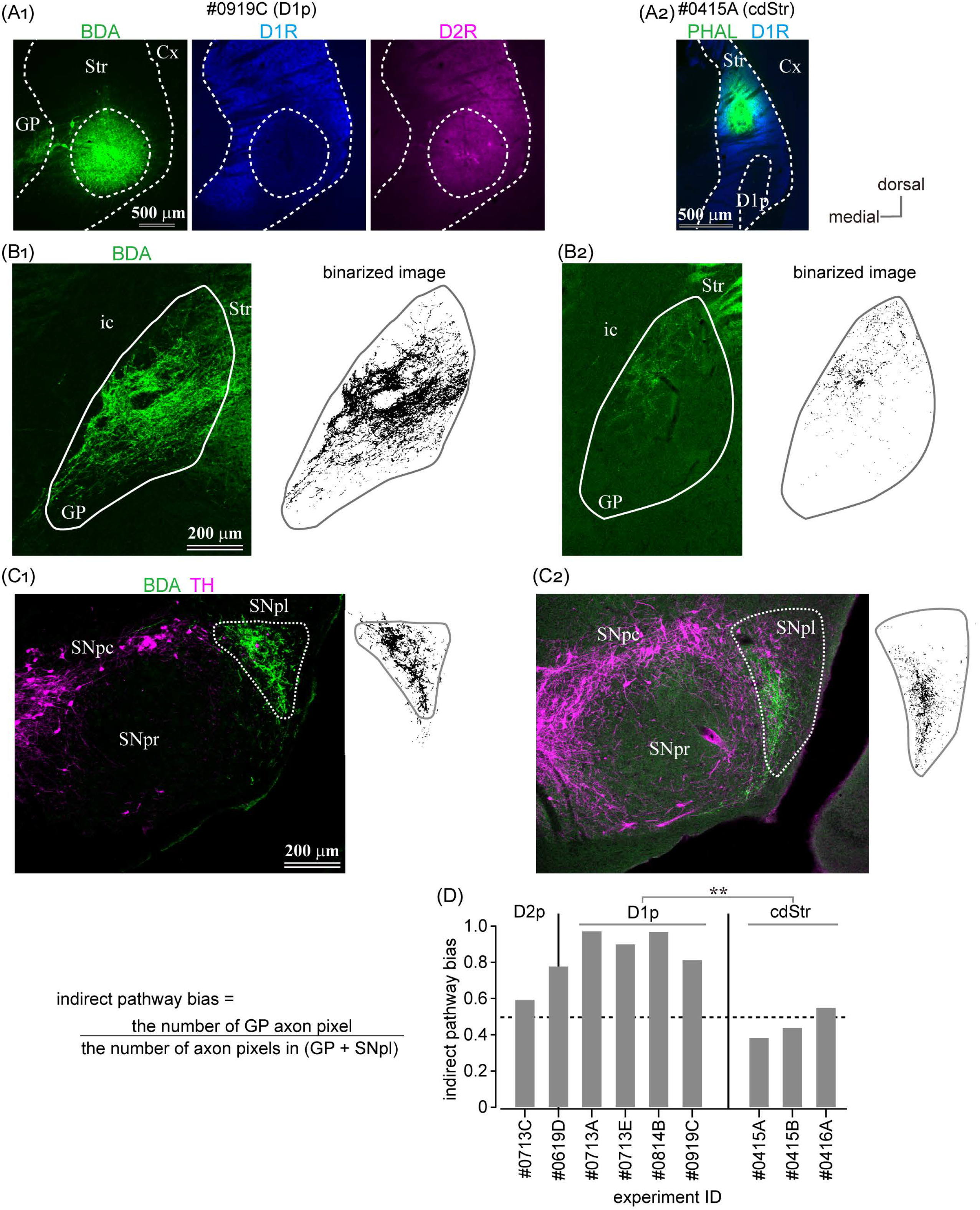
Anterograde axon tracing of the caudal striatum. (A) Representative examples of tracer injection into D1R-poor zone (A1) or the dorsal part of caudal striatum (cdStr) (A2). Biotinylated dextran amine (BDA; green) was injected in D1R-poor zone, whereas Phaseolus Vulgaris Leucoagglutinin (PHAL; green) was injected to cdStr. (B) Photoimages (left) and their binarized plots (right) of labeled axons in GP from D1R-poor zone (B1) or from cdStr (B2). The binarized pixel plots were used for quantification. (C) Photoimages (left) and their binarized plots (right) of labeled axons in SN from D1R-poor zone (C1) or from cdStr (C2). Note dense axons were highly restricted in SNpl. Only few axons in SNpr or SNpc. (D) Indirect pathway bias (see the main text) of each injection. Injection into D1R-poor zone resulted in significantly higher values than that into cdStr. Cx, cerebral cortex; GP, globus pallidus; ic, internal capsule; Str, striatum; SNpc, substantia nigra pars compacta; SNpl, substantia nigra pars lateralis; SNpr, substantia nigra pars reticulata.

**Figure 7.**
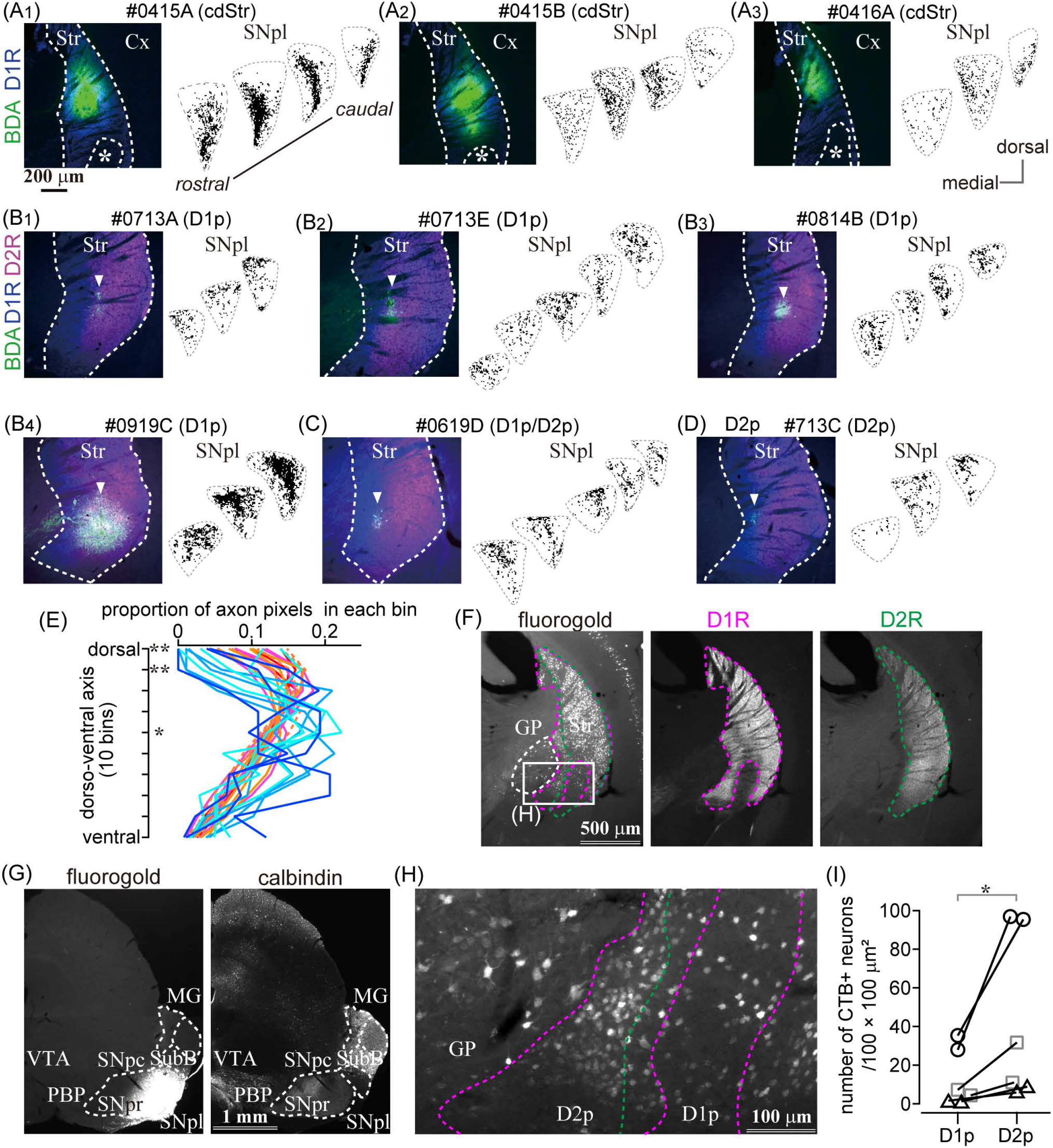
The caudal striatum projects to SNpl. (A)-(D) Anterograde tracer injection sites in the caudal striatum (left) and the binarized axon plots in SNpl (right) for all samples. SNpl drawings are arranged in a rostro-caudal order. (A) Injections into cdStr (N = 3 mice). Asterisk indicates D1R-poor zone. (B) Injections into D1R-poor zone (D1p) (N = 4). (C) Injection into the border between D1p and D2R-poor zone (D2p) (N = 1). (D) Injection into D2p (N =1). (E) Comparison of axon distribution in SNpl along a dorso-ventral axis. In dorsal part of SNpl, axons of D1R-poor zone (reddish lines) were relatively dense than those of cdStr (bluish lines). *, *p* <0.05; **, *p* <0.01. (F)-(I) Retrograde tracer injection into SNpl labeled cStr neurons. (F) Left, Neurons labeled by fluorogold injected into SNpl. Immunostaining for D1R (middle) and D2R (right) is used to determine the poor zone. (G) Fluorogold injection into SNpl (left). The section was counter stained with calbindin-D-28k (right). Fluorogold extended to the lateral part of SNpr. (H) A magnified image of the rectangle area in F. Labeled neurons were located in GP, D1p, and D2p. (I) The number of SNpl projecting neurons in D1R- and D2R-poor zone. Larger number of neurons were labeled in D2R-poor zone. Individual mice were marked with a different symbol (N = 3 mice, two sections/mouse). Cx, cerebral cortex; MG, medial geniculate nucleus; PBP, parabrachial pigmented nucleus; Str, striatum; SNpc, substantia nigra pars compacta; SNpl, substantia nigra pars lateralis; SNpr, substantia nigra pars reticulata; SubB, subbrachial nucleus; VTA, ventral tegmental area.

We also observed the axons in zona incerta, thalamic ventral posterior nucleus (VPM and VPL), thalamic reticular nucleus (Rt), and medial geniculate nucleus (MG). Because in the cStr (including the poor zones), there are several neural fibers are passing through; BDA can lead to ectopic labeling– other than the injection center, through such fibers or terminals in the striatum. We found a few labeled neurons in multiple cortical areas, thalamus, and brainstem probably due to retrograde transport of BDA; thus, the above ectopic labeling may be contaminated with actual striatal projections.

To confirm projection from the poor zones, retrograde tracers, fluorogold or CTB555, was injected into either MG or SNpl (Figure 7F, 7G, Supplementary figure 4). We found prominent retrograde labeling in cStr including both D1R- and D2R-poor zones by SNpl injection, as well as labeling of large sized neurons in the caudal GP (Figure 7F, 7H). The total number of labeled neurons was 249 in D1R-poor zone and 842 in D2R-poor zone (N = 3 mice; two sections/mouse). The density of labeled neurons was significantly larger in D2R-poor zone than in D1R-poor zone (Figure 7I, *p* = 0.031 by paired t-test); this was similar to the case with large volume retrograde tracer injection as shown in Figure 5. On average, the mean ratio of labeled neuron density in D2R-poor zone *vs*. that in D1R-poor zone was 4.05 ± 1.74 (range: 2.51 – 7.28), reflecting the number of the dMSNs in each zone. This result supported the aforementioned anterograde tracing data which showed that both poor zones projected to SNpl. Meanwhile, little to no labeling in cStr was observed for MG injection; rather MG injection provided intensive labeling in the multiple cortical areas—the primary and secondary somatosensory area (S1 and S2); dorsal and ventral auditory area (Au), temporal association area (TEA), and amygdala. Thus, the main target of the direct pathway in the poor zones must be SNpl (see also Supplementary Figure 4).

### 3.6 Uneven distribution of immunoreactivity of D1R and D2R in the caudal striatum across age, sex, strain, and species

Since cStr is innervated by the sensory cortices, such as visual and auditory areas, it may be possible that the presence of poor zones depends on intrinsic factors, such as age and sex. Particularly, it is well-known that aged BL6 mice lose their hearing. We examined the poor zones in young (4-week-old; N = 3; Figure 8A) and aged (N = 3; 61-, 111-, and 172-week-old; Figure 8B) mice. As shown in Figure 8, the poor zones were obviously present irrespective of age. Moreover, even in 8-day-old (P8) mice, the uneven distributions of D1R and D2R were clearly observed (Figure 8C; N = 3 mice).Furthermore, the poor zones were also present in female mice (Figure 8D; N = 3 mice, 10–12-week-old). The line profiles of D1R and D2R immunofluorescence (Figure 8A-D) demonstrated similar spatial distributions to those observed in adult male mice as shown in Figures 1 and 2.

**Figure 8.**
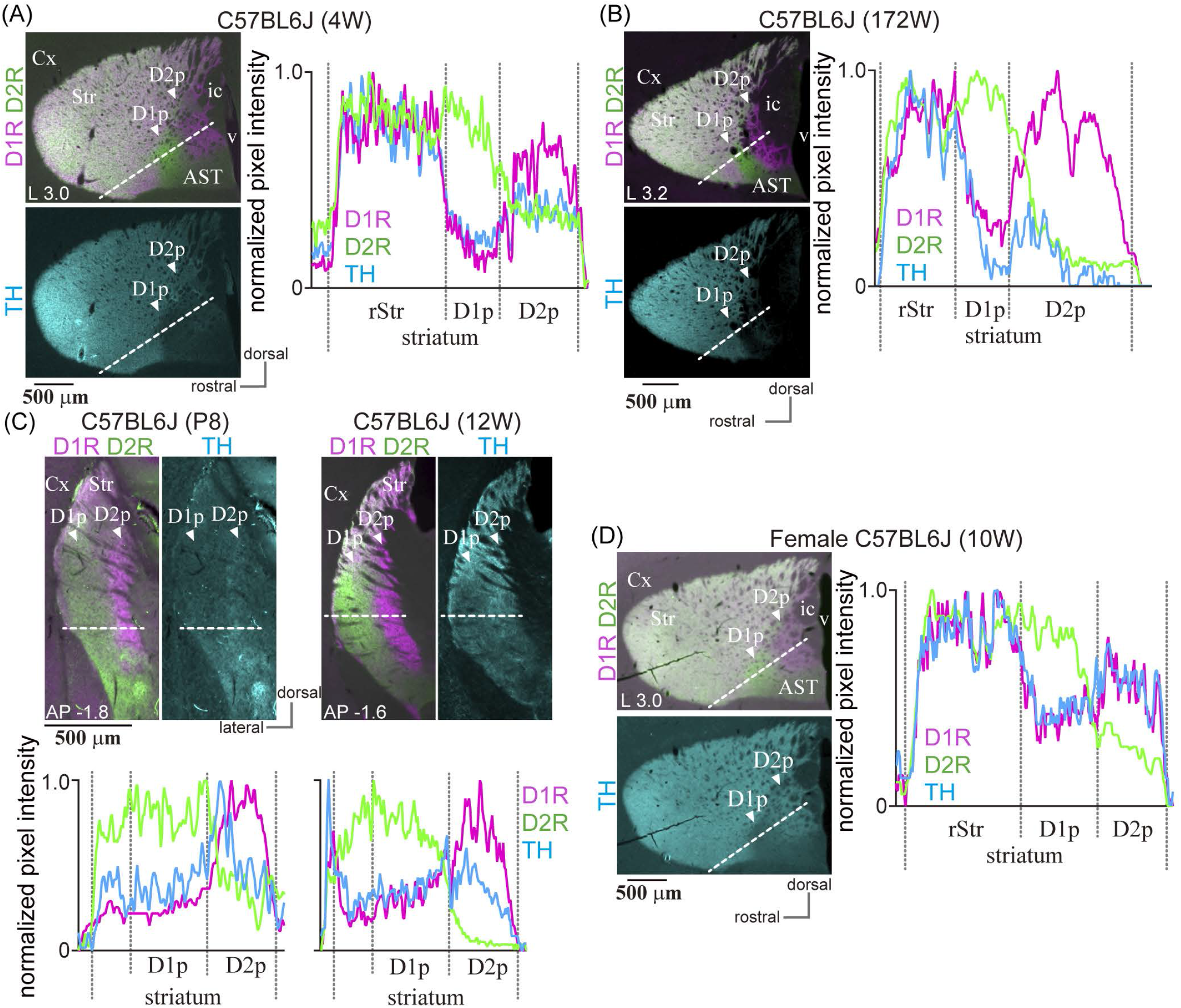
Presence of the poor zones in BL6 mice independent of age and sex. (A) The poor zones in a 4-week-old (4W) male BL6 mouse. The merged image of immunofluorescence against D1R (magenta) and D2R (green) of a L3.0 sagittal section is shown in the left upper panel. The left lower panel represents TH immunofluorescence (cyan) in the same section. The line profiles of fluorescence intensity (right) were measured on the dotted white line shown in the left images. (B) The poor zones in a 172-week-old aged BL6 mouse. A sagittal section at L3.2 is shown. (C) The poor zones already existed in a postnatal 8-day (P8) BL6 mouse. Due to small size of a brain, P8 sections were derived only as coronal sections to obtain enough sections containing the poor zones for multiple series of immunostaining. A coronal plane at AP −1.8 is shown on the left. For comparison, a coronal plane at AP −1.6 of 12-week-old BL6 is shown on the right. The corresponding line profiles are represented in the bottom. (D) Poor zone in a female BL6 mouse (10-week-old) in a sagittal section. Line profiles for D1R and D2R immunofluorescence confirmed a similar pattern of the poor zones irrespective of age and sex.

To determine whether the uneven distribution of immunoreactivity of D1R and D2R in cStr is conserved across strains and species, ICR mice (11-week-old, N = 3), and Wistar and Long-Evans rats (N = 3 of each; 11–13-week-old) were used to test for triple immunofluorescence of D1R, D2R, and TH (N = 3 of each type of animal). Uneven distribution of D1R and D2R was also found in the cStr of ICR mice (Figure 9A), Wistar rats (Figure 9B), and Long-Evans rats (N = 3, Figure 9C), as reported for Sprague-Dawley rats previously (Gangarossa et al., 2019). Similar to that of mice, D2R-poor zone of rats was always located at the most caudal part of the striatum, and D1R-poor zone was located rostral to and medial to D2R-poor zone, although the appearance of these zones (shape and size) was slightly different than those in mice. The boundary of D1R- and D2R-poor zones extended more dorsally in Long-Evans rats than Wister rats (Figure 9B, 9C). In addition, D2R-poor zone spread toward the dorsal edge of the cStr in both strains of rats.

**Figure 9.**
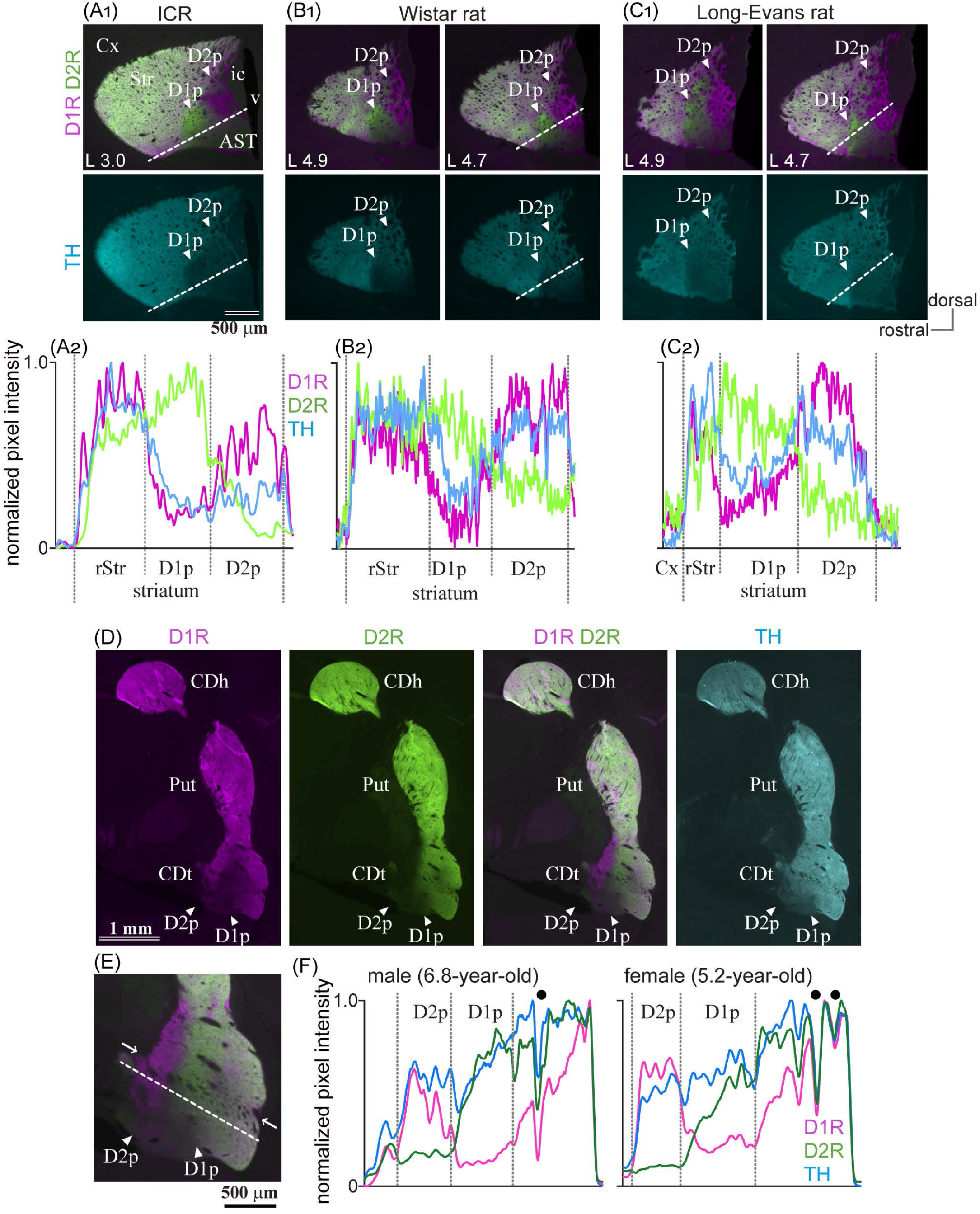
Presence of the poor zones in ICR mouse, rats, and marmosets. (A1)-(C1) Images of immunofluorescent labeling for D1R (magenta), D2R (green), and TH (cyan) in the striatum of rodents. Arrowheads indicate D1R- or D2R-poor zone (D1p or D2p). The distance from the midline is shown at the lower left corner. (A2)-(C2) Line profiles of pixel intensity for immunofluorescence of D1R and D2R in ICR mouse and rats. The dotted white line for quantification of the pixel intensity is shown in the top panel. (D) Uneven distribution of D1R D2R, and TH in the caudate of marmosets (coronal plane at ∼ AP 3.3). (E) A magnified view of the CDt, including the poor zones (arrowheads). Note nerve bundles (arrows) crossing the striatum containing the poor zones. (F) Line profiles of the marmoset. Filled black circles represent location of nerve bundles or large blood vessel. AST, amygdala striatal transition area; CDh, caudate head; CDt, caudate tail; Cx, cerebral cortex; ic, internal capsule; v, ventricle.

To determine whether the uneven distribution of D1R, D2R, and TH is specific to rodents, the striatum of common marmosets—a nonhuman primate, was also examined (N = 1 male and 1 female, Figure 9D, 9E, 9F). In primates, the striatum exclusively subdivides further into two nuclei, the putamen and caudate nucleus. In the marmoset, the unique region with uneven distribution of D1R and D2R was found around the rostral tip of tail of the caudate (CDt). D1R- and D2R-poor zones were adjoined, and D1R-poor zone was located laterally to D2R-poor zone, as observed in rodents. This uneven distribution zones spread beyond thick nerve bundles (arrows in Figure 9E), which crosses the striatum, and is considered as the border between the putamen and CDt. However, it is difficult to determine whether the spatial composition in the marmosets is similar to that in the rodents owing to anatomical structural differences of the respective striata.

## 4 Discussion

The present study demonstrated that the rule of cellular correlation between dopamine receptor expression and projection pathways was conserved in the unusual part of the cStr where D1R-MSNs and D2R-MSNs are unevenly distributed. To the best of our knowledge, this is the first report which describes axonal projections from D1R-poor zones, although Gangarossa et al. (2013b) have reported that D2R-poor zone projected to SNpr and EP (see also Valjent & Gangarossa, 2020). We identified the characteristic projections from the poor zones to SNpl via the direct pathway and to the caudal GP via the indirect pathway (Figures 6, 7, and Supplementary Figure 2, 3, 4). We have also elucidated that the electrophysiological properties of MSNs in the poor zones were similar to those observed in other striatal area for most of the parameters evaluated in this study (Figures 4, Table 3), and identified the poor zones in primates (Figure 9).

### 4.1 Methodological consideration

The present axonal tracing method may be considered to be limited, as the tracer deposit in the poor zones could be extremely small, causing incomplete labeling. Therefore, we cannot exclude the possibility of additional projections to other brain areas, not detected in the present study, or of potential differences in projections from two poor zones. However, even then, the projections that originated from a smaller population of MSNs in each poor zone, namely, the direct pathway in D1R-poor zone or the indirect pathway in D2R-poor zone, were successfully visualized. Thus, it is highly likely that characteristics of principal projections from the poor zones were correctly obtained, irrespective of the technical limitations. In addition, it should be mentioned that our electrophysiological experiments were also limited in regards to the sample size and the parameters analyzed. The present results simply showed that most of the basic membrane properties of MSNs in the poor zones were not significantly different from that of cdStr MSNs using *in vitro* preparations. However, as a significant difference was observed in the rheobase current, it could be also possible that there was some differentiation of detailed electrophysiological properties of MSNs in the poor zone. The differences of dopaminergic innervation and composition of interneurons (Miyamoto et al., 2019) can affect synaptic integration and plasticity. In addition, the axonal projections revealed by the microinjections of anterograde tracers were mainly derived from D1R-poor zone. The possibility of differences in projections from the neurons in D1R- *versus* D2R-poor zones cannot be excluded, although SNpl was a target for projections from both poor zones (Figure 7).

### 4.2 Dual pathway neurons in D1R- and D2R-poor zones

In transgenic mice, uneven distribution of D1R and D2R expressing neurons in the cStr was reported (Gangarossa et al., 2013b, 2019; Miyamoto et al., 2019); the present study used wild type animals including C57BL/6J mice and our findings of the locations of the unique zones and of cell type composition were consistent with these previous reports (Figures 1, 2A–2C).

Cellular distribution is not always precisely the same as distribution of immunoreactivity, because the protein—visualized by immunohistochemistry, is localized at not only the soma but also in the dendrites and axons. To account for this, we performed *in situ* hybridization study and revealed the complementary distribution of Drd1a and Drd2 expressing neurons in both D1R- and D2R-poor zones (Figure 3H). These findings suggest the possibility that the direct and indirect pathway neurons are unevenly distributed in the cvStr.

To address this question of uneven distribution of projection neurons, we initially performed a retrograde tracing study combined with immunohistochemistry. The proportion of retrogradely labeled striatonigral neurons, considered as direct pathway neurons, were significantly lower in D1R-poor zone and significantly higher in D2R-poor zone than other regions (Figure 5A–5D). It is noteworthy that the estimated density of striatal neurons as observed by our large volume tracer injections is consistent with an early report (Rosen & Williams, 2001), suggesting that a large population of direct pathway neurons can be labeled and the differences in the labeled neuron density are not caused by biased topographic distribution of projection neurons. In addition, the combined study of *in situ* hybridization with retrograde labeling showed that the retrogradely-labeled direct pathway neurons expressed Drd1a, but not Drd2 mRNA (Figure 3E, 3F). These findings indicated that the distribution of direct pathway neurons in the cvStr is highly biased toward D2R-poor zone. Owing to the existence of axon collaterals of the direct pathway neurons in GP (Fujiyama et al., 2011; Kawaguchi et al., 1990; Levesque and Parent, 2005; Wu et al., 2000), it is difficult to label only the indirect pathway neurons using neural tracers. Nevertheless, our data suggested that D2R-neurons in the poor zones are likely to target GP as those in other striatal areas, because the D1R-poor zone projection was denser in GP than in EP and/or SN (Figures 6, 7, and Supplementary Figure 2). In addition, retrograde tracer injection revealed that both caudal GP and cStr convergently projected to SNpl (Figure 7F–7H). Anterograde tracing also suggested that the caudal GP is likely to constitute the indirect pathway of cStr including poor zones (Figure 6, Supplementary Figure 2), as reported in primates (Amita and Hikosaka, 2019). Thus, the direct and indirect pathway neurons in the poor zones could form complementary circuitry, even though they were distributed unevenly. An assumption of the potential relationship between the poor zones and the striosome (patch)/matrix compartments may be implicated. However, in cvStr, typical striosome/matrix compartmentalization disappears (Miyamoto et al., 2018, 2019).

We also confirmed the preservation of D1R- and D2R-poor zones across age and species (Figures 8, 9); however, we did not conduct any experiments on their functional relevance. Thus, the necessity and/or significance of such unique regions is still unknown. In marmosets, the poor zones are likely to contain the ventral end of the putamen, the border region of putamen/CDt, as well as the rostral end of CDt. Alternatively, at least in this part of the striatum, the extent of CDt itself may spread out more dorsally than that of more caudal CDt which did not contain the poor zones. A recent on-line atlas on marmosets (https://gene-atlas.brainminds.riken.jp; Kita et al., 2021; Shimogori et al., 2018) supported the uneven distribution of D1R/D2R expressing cells (Drd1, ID#369-1 and #N-2; Drd2, ID #AA-2). The cStr, including poor zones, is innervated by the sensory cortices, which implies a possible relationship between sensory inputs and the poor zone development. However, this may not be the case. The poor zones were already present at ∼P8, when behavioral reactions to tones or to visions were not yet elicited (Ehret, 1976; Huberman et al., 2008). Moreover the poor zones were preserved in aged C57BL6/J mice even though they were generally hard of hearing (Zheng et al., 1999); however, sensory deprivation affects synaptic transmission in the sensory striatum (Mowery et al., 2017). Thus, structure of the poor zones could develop and be maintained without sensory signals.

### 4.3 The caudal striatum and the poor zone in rodents and primate

Recent studies with primates and rodents throw light on the functional aspect of the cStr. In the macaque monkey, the CDt, which is a long C-shaped structure of the ventral part of the caudate nucleus, has been reportedly involved in the distinct value coding (Griggs et al., 2017; Kim et al., 2017; Kim et al., 2014; Kim and Hikosaka, 2013, 2015). These findings can also be explained by the CDt receiving the distinct subpopulation of dopaminergic neurons in SNpc. Meanwhile, the poor zones in the common marmosets revealed here were located in approximately the most rostral end of the CDt. This part of the CDt is small; therefore, its neural connections and functions are still not clear. Contrastingly, Menegas et al. (2017, 2018) reported that the tail of the striatum in rodents is involved in saliency and aversion. In addition, the shape of the poor zones differed even in mice and rats (Figure 8). Thus, anatomical and functional similarity of the poor zones and the tail of the striatum among species warrants further research.

### 4.4 Functional implications of the poor zone

We have reported that the main projection target of the cStr direct pathway, including the poor zones, was SNpl. The previous studies reported that cStr subregions receiving auditory cortical inputs project to the SNpl, whereas cStr subregions receiving visual cortical inputs project to the lateroventral SNpr (Deniau et al., 1996; Kohno et al., 1984). Such meso-scale topographic relationship was also reported in primates (Hedreen and DeLong, 1991). In turn, ventral SNpl is known to project to the superior colliculus (SC), inferior colliculus (IC), VPM, and so on; particularly, it innervates the sensory-related regions of the thalamus and midbrain (Cebrian et al., 2005; Moriizumi et al., 1992; Takada, 1992; Yasui et al., 1991). A recent study showed that SNpl was densely innervated by GABAergic neurons in the central amygdala (see also Supplementary Figure 4B), and the connection could relate to appetitive and aversive learning (Steinberg et al., 2020). Thus, the central amygdala and cStr have common characteristics on projections to SNpl, and on functional relevance to saliency and aversion. In addition, cStr is innervated by sensory related cortical areas: auditory, visual, secondary sensory, TeA, and perirhinal areas (Hintiryan et al., 2016; Hunnicutt et al., 2016; Jiang and Kim, 2018; Yeterian and Pandya, 1998; Yeterian and Van Hoesen, 1978). Recently, Li et al. (2021) reported that crescendo auditory stimuli, which mimic an approaching subject, induced freeze and flight responses in mice. Interestingly, corticostriatal innervation from the auditory cortex preferentially innervated D2R- neurons, in turn, excitation of these D2R-neurons increased freezing, although the precise relationship between the neural/behavioral responses and the poor zones are not reported. Taken together, cStr can contribute to the gating and integration of multimodalities of sensation via SN in rodents, similar to SN-subregion dependent information coding reported in primates (Amita et al., 2020; Kim and Hikosaka, 2015; Yamamoto et al., 2012; Yasuda and Hikosaka, 2015). However, it may be noteworthy that most of neural activity data in primates have been obtained from a more caudal part of CDt. Our anterograde and retrograde tracing also suggested that the caudal GP is likely to constitute the indirect pathway of cStr including poor zones, which projected to SNpl (Figures 6, 7), as reported in primates (Amita and Hikosaka, 2019). Furthermore, cdStr and poor zones tended to project slightly different parts of the SNpl (Figure 7A–7E, Supplementary Figure 3), which suggests a potential functional differentiation between cdStr and the poor zones. The topographic projection from cStr has been previously reported (Foster et al., 2021), although the poor zones were not considered in this paper.

### 4.5 Is the poor zone an exception of the striatum?

The classical model for the basal ganglia network suggests that normal function requires a balance of the direct and indirect pathway neurons mediated by D1R and D2R, respectively. The uneven distribution of the two MSNs population in the cStr raised a question regarding the balance between the direct and indirect pathway neurons. If the balance in the poor zones was as critical as other striatal areas, the two poor zones would share neural information with the aid of common inputs and/or mutual connections, or local circuitry within each zone, although our neural tracing experiments did not provide concrete evidence for these possibilities. Precise and detailed neural circuitry in these zones should be explored in future research. Whether D1R- and D2R-poor zones share common cortical inputs remains to be determined; if they do, two poor zones can work as one unit, like other striatal area containing equal number of D1R- and D2R-neurons. It is also possible that both poor zones could be innervated by different populations of cortical neurons. In other striatal regions, the cortical axons in the adjoining striatal areas originated from the adjacent but segregated cortical regions (Ghosh and Zador, 2021; Hooks et al., 2018). In such a case, two poor zones might communicate to compensate for the highly biased distribution of the direct and indirect pathway neurons, if they still need to work in a coordinated manner.

Alternatively, unlike motor- or limbic-related information processing in the striatum, these poor zones could work via either the direct or indirect pathway. This view is supported by a recent research that showed highly biased corticostriatal innervation onto D2R-neurons from the auditory cortex (Li et al., 2021). In such an event, lesser dopaminergic innervation specifically observed in the poor zones, which is likely to be from the SNpl (Jiang and Kim, 2018; Menegas et al., 2015; Poulin et al., 2018; Watabe-Uchida et al., 2012) could also relate to such unusual striatal circuitry. Because it is also suggested that the nucleus accumbens, a part of the limbic system, also contains a poor zone-like structure (Gangarossa et al., 2013a; Petryszyn et al., 2017), the unique circuit may not only relate to the sensory system. Furthermore, to determine whether the poor zones and cdStr are functionally differentiated, and whether the cStr is a counterpart of the primate CDt, neural tracing and functional recording/imaging in extremely fine scales is required, which will help to understand the two pathways beyond the current concept.

## Abbreviations

ACSF: artificial cerebrospinal fluid
AP: antero-posterior
AST: amygdala-striatal transition area
Au: auditory area
BDA: biotinylated dextran amine
CDt: caudate tail
cStr: caudal striatum
cdStr: caudo- dorsal striatum
CTB: cholera toxin subunit B
DAT: dopamine transporter
dStr: dorsal striatum
D1R: dopamine receptor D1
D2R: dopamine receptor D2
EP: entopeduncular nucleus
FG: fluorogold
GP: globus pallidus
LM: latero-medial
MG: medial geniculate nucleus
MSN: medium spiny neuron
PB: phosphate buffer
PBS: phosphate buffered saline
PHAL: phaseolus vulgaris leucoagglutinin
ppz: para-poor zone
rStr: rostral striatum
RT: room temperature
SNpc: substantia nigra pars compacta
SNpl: substantia nigra pars lateralis
SNpr: substantia nigra par reticulata
SubB: subbrachial nucleus
TH: tyrosine hydroxylase
VPM: ventral posterior nucleus of thalamus, the medial part
VPL: ventral posterior nucleus of thalamus, the lateral part

## 6 Author Contributions

All authors had full access to the data in the study and take responsibility for the integrity of the data and the accuracy of the data analysis. Conceptualization, F.F. and F. Karube; Methodology, F. Karube, Y.H., and K.O.; Investigation, K.O., F. Karube, F. Kadono, Y.H., and F.F.; Formal Analysis, K.O., F. Kadono, F. Karube, and Y.H.; Resources, K.I. and M.T.; Writing - Original Draft, K.O., F.F., and F. Karube; Writing - Review & Editing, all authors; Visualization, K.O., F. Karube; Supervision, F.F. and F. Karube; Funding Acquisition, F.F. and F. Karube.

## Acknowledgments

We thank Dr. H. Hioki and Dr. S. Okamoto for their suggestions on *in situ* hybridization. We thank Dr. S. Takahashi and Mr. T. Higashiyama for their helpful insights. We also extend our gratitude to Dr. Y. Sakurai for supervising the study and for his kind encouragement.

## Supplementary Material

### Supplementary Figures

**Supplementary Figure 1.**
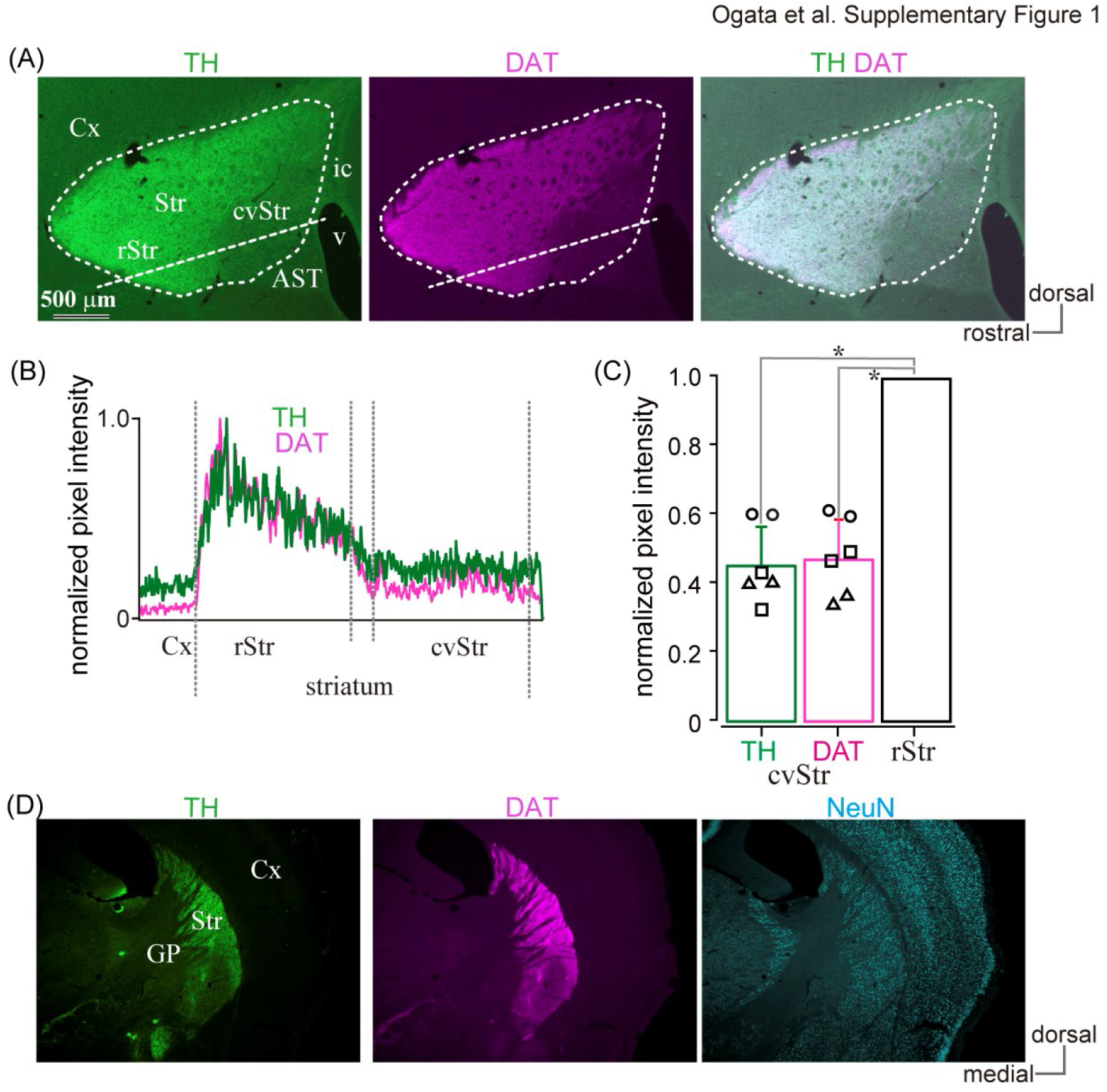
Co-expression of tyrosine hydroxylase (TH) and dopamine transporter (DAT) in the caudal striatum. (A) Immunofluorescent images against TH and DAT in the mouse striatum. Expression of both TH and DAT was low in the caudoventral part of the striatum (cvStr) than in the rostral striatum (rStr) in sagittal plane. (B) Line profiles of TH (green) and DAT (magenta) along with straight dotted lines shown in A. TH and DAT profiles correlated highly. (C) Normalized pixel intensity of TH and DAT was significantly lower in the poor zone (N = 3 mice, each mouse was represented with a different mark; 2 ROIs/section, one section/mouse). (D) Coronal plane. AST, amygdala striatal transition area; Cx, cerebral cortex; GP, globus pallidus; ic, internal capsule; v, ventricle.

**Supplementary Figure 2.**
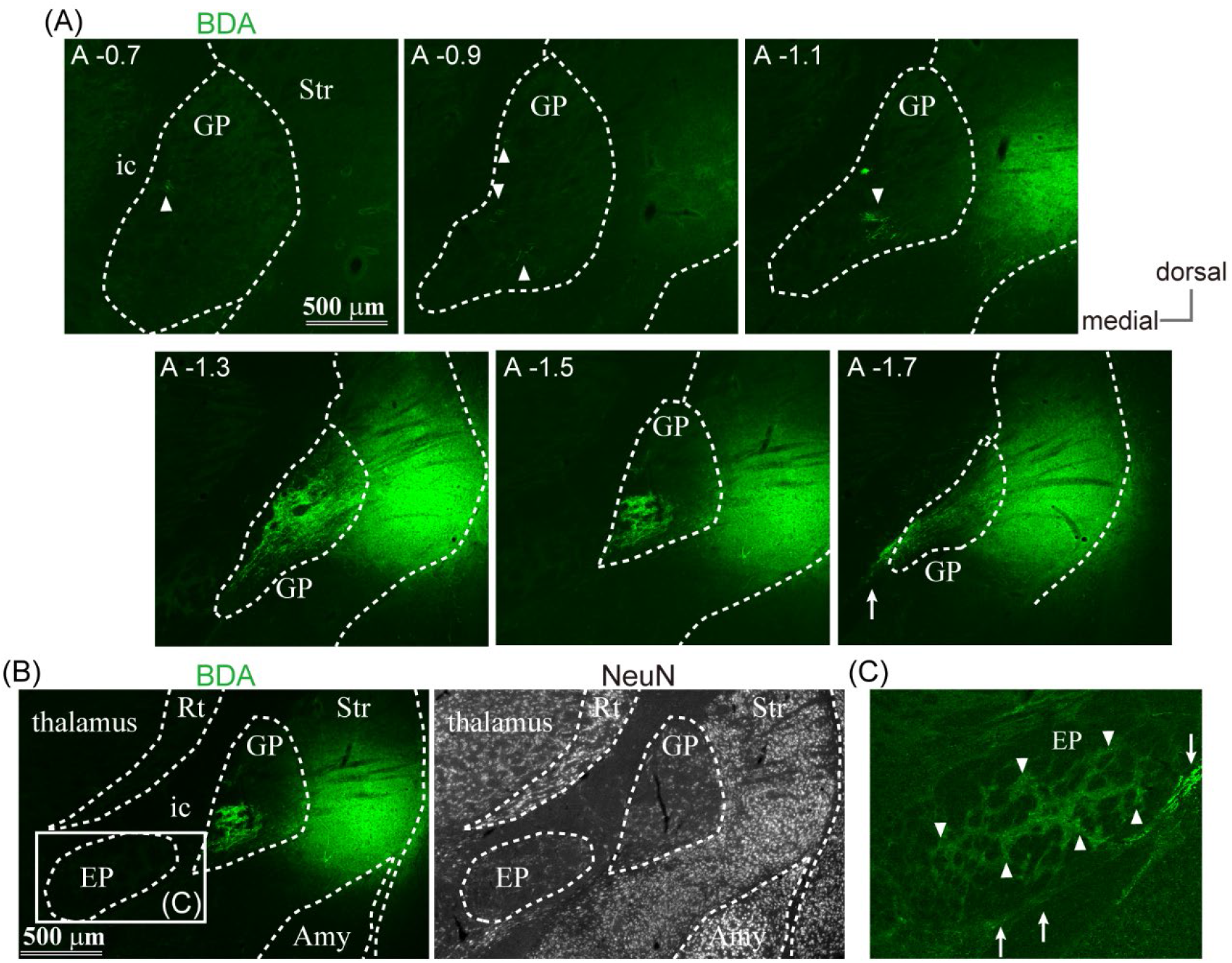
Anterogradely labeled axons in globus pallidus (GP) and entopeduncular nucleus (EP) from D1R-poor zone. (A) For biotinylated dextran amine (BDA) injection to poor zones, labeled axons were found in the caudal globus pallidus (GP), and only few axons were found in the rostral GP (arrowheads). Arrow indicates descending axons toward the downstream nuclei. The images were arranged from rostral (the upper left) to caudal (the lower right). (B), (C) Only few axons were labeled by injection to cStr in EP (cdStr and the poor zones). (B) Left, BDA labeled axons in low magnification. Right, NeuN immunostaining. (C) A magnified view of rectangle area shown in B. Apart from descending axons passing internal capsule (arrows), there were few axon in EP (arrowheads).

**Supplementary Figure 3.**
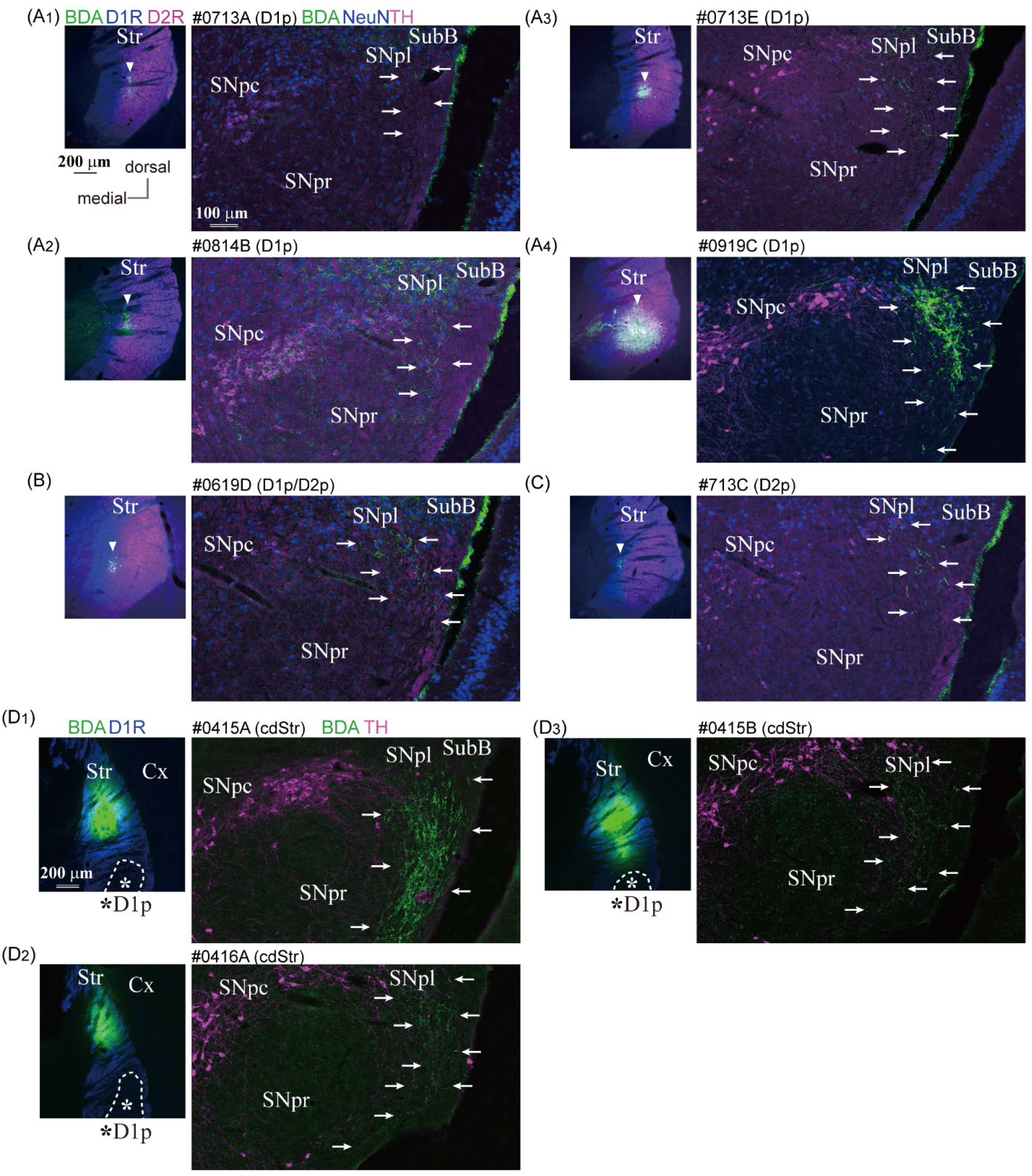
Projection from the poor zones and the dorsal part of caudal striatum (cdStr) to substantia nigra pars lateralis (SNpl). (A) Micro injection of biotinylated dextran amine (BDA) to D1R-poor zone in order of the size of injection core (A1–A4). Left, The core of BDA injection (green; arrowhead) with immunofluorescent labeling for D1R (blue) and D2R (magenta). The core is located in D1R-poor zone. Right, BDA-labeled axons were restricted in SNpl (arrows). (B) Injection to the border between D1R-poor zone and D2R- poor zone. (C) Injection to D2R-poor zone. In all the cases, most of the axons were restricted in the dorsal part of SNpl, and only few axons were present in SNpc or SNpr. (D) Injection of Phaseolus Vulgaris Leucoagglutinin (PHAL) in the dorsal part of caudal striatum (cdStr) where both D1R and D2R co-expressed. Axons from cdStr were also found in SNpl, with collaterals into the ventral part of SNpl. SubB, subbrachial nucleus.

**Supplementary Figure 4.**
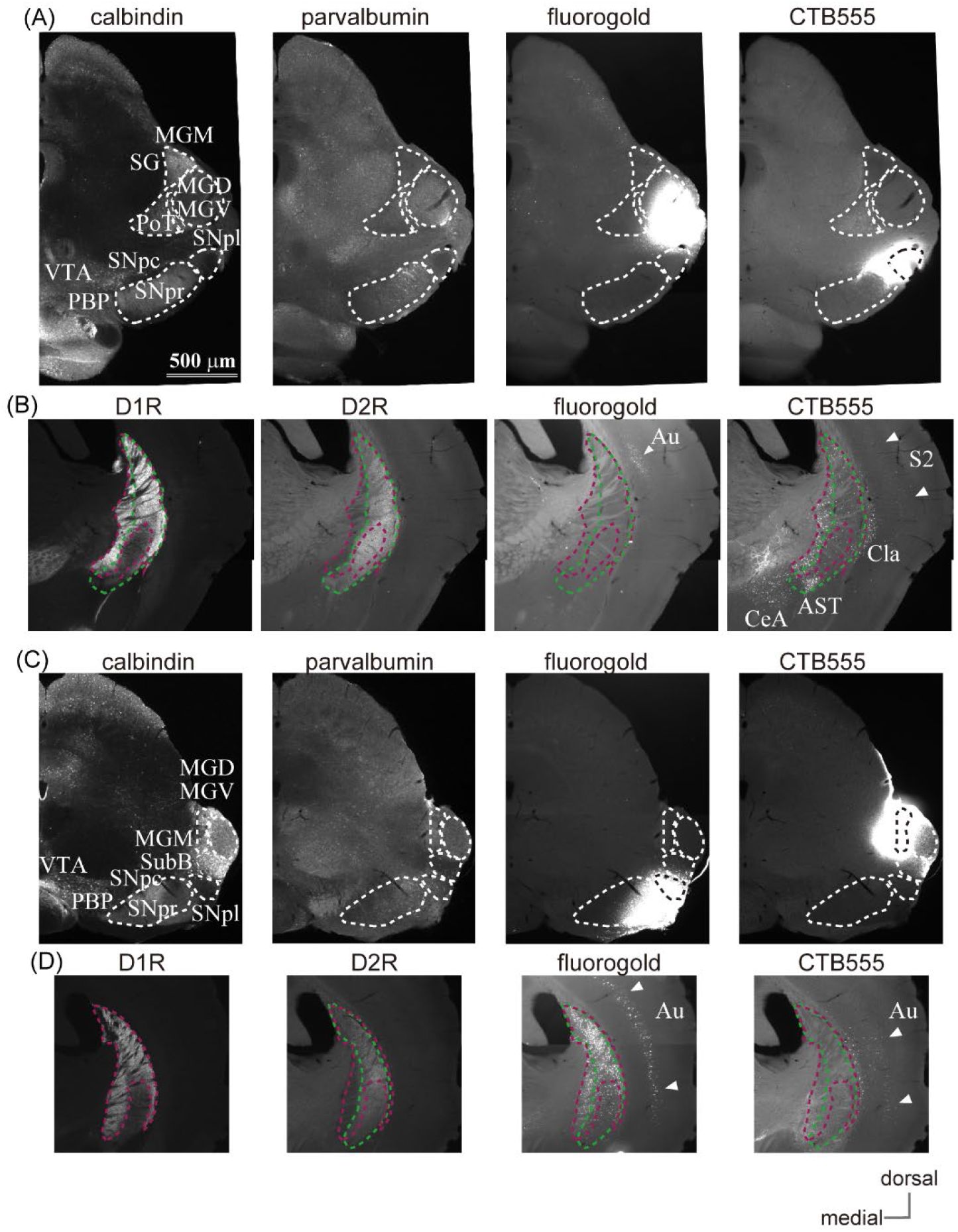
Retrograde tracing confirmation of caudal striatum (cStr) projection. (A) An example of double retrograde tracer injection into MG and SNpl. Fluorogold was injected into MGV/MGD, whereas CTB555 in the dorsal SNpl. The substructures of the midbrain were determined by the immunofluorescent staining patterns against calbindin and parvalbumin. (B) Distribution of labeled cell bodies. Immunofluorescent labeling for D1R and D2R were used for identification of poor zones in the caudal striatum (cStr). Fluorogold injection into the MG labeled deep layer neurons in temporal cortical area, including auditory related areas (Au, arrowheads). They were also found in other temporal areas (not included in this section). Only few striatal neurons were labeled. Meanwhile, CTB injection into SNpl densely labeled cStr neurons, that were especially numerous in D2R-poor zone. Cortical neurons in deep layers were also labeled (arrowheads; corresponding to the secondary sensory area (S2) in this section). (C) Another example of fluorogold injection into the ventral SNpl and SNpr, and CTB injection into MG. (D) Dense fluorogold labeled neurons in cStr, being relatively dense in the dorsal part of cStr. Again, D2R-poor zone contained more labeled neurons than D1R-poor zone. Labeled neurons were also found in the deep layers of the temporal cortex (arrowheads). CTB injection did not result in labeling of the striatal neurons, rather, deep cortical neurons were labeled. Au, auditory cortex; AST, amygdala striatum transition area; CeA, central amygdala; Cla, claustrum; MGM, medial part of medial geniculate nucleus; MGD/MGV, dorsal/ventral part of medial geniculate nucleus; PBP, parabrachial pigmented nucleus; PoT, triangular part of posterior thalamic nuclear group; S2, the secondary somatosensory cortex; SNpc, substantia nigra pars compacta; SNpr, substantia nigra pars reticulata; SNpl, substantia nigra pars lateralis; SubB, subbrachial nucleus.

